# Prior infection with IBDV prolonged the shedding of a mallard H3N8 influenza A virus (IAV) challenge from the oropharyngeal cavity of some chickens and increased the number of amino acid substitutions in the IAV samples

**DOI:** 10.1101/2024.12.31.630863

**Authors:** Salik Nazki, Chandana Tennakoon, Vishwanatha R. A. P. Reddy, Yana Chen, Jean-Remy Sadeyen, Andrew J. Brodrick, Munir Iqbal, Holly Shelton, Andrew J. Broadbent

**Author notes:** Correspondence: Andrew J. Broadbent.

## Abstract

Infectious bursal disease virus (IBDV) is endemic worldwide and causes immunosuppression in chickens. We hypothesized that a previous history of IBDV in chickens would render them more susceptible to infection by influenza A viruses (IAVs) from aquatic waterfowl reservoirs. To model this, we inoculated 14 day old specific pathogen free (SPF) chickens with a low pathogenicity avian influenza (LPAI) virus strain from a mallard (A/Mallard/Alberta/156/01 (H3N8)) and compared replication and shedding between immunocompetent chickens and chickens that had immune dysregulation due to a prior IBDV infection with strain F52/70 (genogroup A1B1) at 2 days of age. The mallard IAV strain replicated in the upper respiratory tract of the chickens, and virus was shed from the oropharyngeal cavity, but there was no shedding from the cloaca, and no transmission to sentinel chickens. Replication of the mallard IAV in the chicken host was associated with amino acid substitutions in the polymerase complex and HA. IBDV infection increased the average fold change of IAV replication in the trachea of chickens, prolonged the shedding of infectious IAV from 5 to 6 days in some chickens, increased the number of amino acid substitutions detected in the IAV population from 13 to 30, and significantly increased the number of mutations per IAV sample from 2.50 (SD +/- 1.83) in the Mock/IAV group to 4.75 (SD +/- 1.81) in the IBDV/IAV group (p < 0.01). Taken together, IBDV infection prolonged the shedding of the mallard IAV in some chickens and changed IAV intra-host evolution.

**Author summary:** Spillover of IAVs from wild aquatic waterfowl into poultry populations occur frequently, which increases the risk of human infection as people have more contact with poultry than wild birds. Poultry flocks may have other co-morbidities that may influence the spread of IAV. Our data demonstrate that prior IBDV infection increased the average fold change of a mallard H3N8 LPAI virus in the trachea of inoculated chickens, prolonged the shedding of infectious IAV from the oropharyngeal cavity, and significantly increased the average number of amino acid substitutions per IAV sample. We hypothesize that IBDV infection could increase the amount of IAV shed into the environment and broaden the diversity of the IAV population shed. We conclude that controlling the spread of wild aquatic waterfowl strains of IAV in chickens should involve a holistic approach, including the control of co-morbidities and immunosuppressive diseases that could exacerbate their spread.

## Introduction

Wild birds, particularly those associated with wetlands (aquatic waterfowl) are the main reservoir for influenza A viruses (IAV) in nature, which are commonly spread to different geographical regions by migration. The main envelope glycoproteins of IAV are the Hemagglutinin (HA, H) and Neuraminidase (NA, N) proteins, and 16 HA and 9 NA subtypes have been identified in wild birds. Antigenic drift and reassortment are known to occur, and the genetic variation in IAV has resulted in the establishment of genetic lineages that are further divided into sub-lineages and clades (1). Spillover of IAVs from aquatic waterfowl to domestic poultry occurs frequently, and several subtypes of IAV have been identified in poultry populations, for example H2, 3, 4, 5, 6, 7, 9, 10, and 11 (2–10). Once in a poultry flock, IAVs can be transmitted directly between individuals, or indirectly through contaminated water, fomites or air (11). Outbreaks of IAV in poultry are usually transient, but selection of adaptive mutations has led to the establishment of lineages that have become endemic in poultry populations in some countries, for example H9N2 in Asia, the Middle East and parts of North and Central Africa (2). In countries where IAVs are not yet endemic in poultry, a test-and-cull approach is often employed to prevent their spread, which has led to the depopulation of millions of birds in the recent outbreaks of H5 viruses in 2014/15 and 2022-2024 (12).

Given the ongoing threat of spillover of IAVs into poultry populations, it is essential that we understand the factors that drive the transmission of IAVs from aquatic waterfowl to poultry. An inherent risk of raising large homogeneous populations of animals in commercial operations is immunosuppression. While nutritional and environmental factors such as heat stress can compromise the immune status of a flock, the majority of cases of immunosuppression in poultry are caused by immunosuppressive viral infections (13). Among the most common and most studied of these is infectious bursal disease virus (IBDV), which is endemic worldwide in chicken populations (14). IBDV is a highly contagious member of the *Birnaviridae*, causing an acute, lytic infection of avian B cells, the majority of which reside in the bursa of Fabricius (BF). Chickens exposed to IBDV typically respond less well to vaccination programs and are more susceptible to secondary infections (14). The economic impact of IBDV is therefore two-fold: first, due to losses associated with morbidity and mortality, and second through indirect losses due to immunosuppression (15).

A link between IAV and IBDV has already been established in chicken populations: In one study, a 1:1 matched case-control study conducted in 16 districts of Punjab and one administrative unit of Pakistan (with 133 confirmed positive farms matched to 133 negative control farms) identified IBDV as a significant factor associated with H9N2 risk in a multivariable conditional logistic regression model (OR: 3.05, CI: 1.82-5.13, p <0.001) (16). In another study, chickens vaccinated with an inactivated H7 IAV vaccine had lower geometric mean antibody titers when they had previously been exposed to IBDV. Moreover, birds that been exposed to IBDV and then vaccinated and challenged had similar mortality rates to those that had not received an IAV vaccine. The authors concluded that IBDV exposure could contribute to poor IAV vaccine efficacy in the field (17). IBDV has also been shown to exacerbate the pathogenesis and shedding of poultry strains of IAV: In one study, the tracheal lesion score of chickens inoculated with A/chicken/Egypt/FAO-S33/2021 (H9N2) was significantly higher 8 days post challenge (dpc), and shedding from the trachea and cloaca was increased at 4 and 8 dpc in birds that had previously been vaccinated with a live IBDV vaccine, compared to immunocompetent birds that had not received the live IBDV vaccine (18). In another study, the genome copy number of IAV strain A/chicken/Iran/772/1998 (H9N2) was higher in the trachea at day 1 post-inoculation (dpi) in chickens that had previously been infected with IBDV, compared to immunocompetent birds, although this was not observed at other time points (19). Furthermore, in a study conducted in turkeys, IBDV inoculation significantly increased the replication of IAV A/chicken/Iran/688/1999 (H9N2) in the trachea and lungs, as determined by RTqPCR (20).

Finally, it was possible to adapt a mallard IAV strain to chickens by serial passage in birds that had previously been inoculated with IBDV, but not in birds that lacked IBDV exposure: The A/Mallard/Pennsylvania/10218/1984 H5N2 virus did not replicate in chickens that lacked IBDV exposure, but was able to replicate in 2/3 (67%) of chickens that had recovered from an IBDV inoculation. This permitted the serial passage and adaption of the IAV to the chickens, with the adapted virus showing an increase in pathology and viral replication in multiple organs compared to the parental strain (21).

Based on these reports, we hypothesized that a previous history of IBDV infection in chickens would render them more susceptible to infection by IAV from an aquatic waterfowl reservoir, allowing for an enhanced potential for introduction of IAVs into chicken flocks. To address this, we provide data on how IBDV infection affected the replication, shedding, and evolution of an aquatic waterfowl IAV challenge in chickens.

## Results

### IBDV strain F52/70 caused immune dysregulation in chickens that were subsequently challenged with the mallard H3N8 IAV

Chickens belonged to the Rode Island Red (RIR) line and were specific pathogen free (SPF). Chickens were divided into 6 groups: Group 1: Mock/Mock (n = 12), Group 2: IBDV/Mock (n = 12), Group 3: Mock/IAV (n = 12) + mock sentinels (S^mock^) (n = 6), Group 4: Mock/IAV (n = 12) + IBDV-infected sentinels (S^IBDV^) (n = 6), Group 5: IBDV/IAV (n = 12) + S^mock^ (n = 6), and Group 6: IBDV/IAV (n = 12) + S^IBDV^(n = 6). Birds were inoculated with IBDV strain F52/70 at 2 days of age, and then IAV strain H3N8 14 days later (Fig 1). To confirm the IBDV inoculation led to immune dysregulation in birds that were subsequently infected with the mallard H3N8 IAV, one day prior to IAV inoculation, a blood sample was obtained from 6 birds per group, PBMCs were isolated, and stained with a cocktail of anti-Bu1-FITC, anti-KUL01-PE, anti-CD4-PE/Cy7, anti-CD8α-Pacific Blue, and Live/Dead™ fixable Near-IR dead cell stain to quantify the percentage of B cells, monocytes/macrophages, CD4^+^ T cells, and CD8^+^ T cells present (Fig 2). Samples from the mock-inoculated birds were compared to samples from IBDV-infected birds. There was a significant reduction in the average percentage of B cells (Fig 2A), and a significant increase in the average percentage of monocytes/macrophages and CD4^-^CD8^+^ T cells (Fig 2B and D), demonstrating that IBDV induced an immune dysregulation in inoculated chickens one day prior to IAV inoculation.

**Fig 1.**
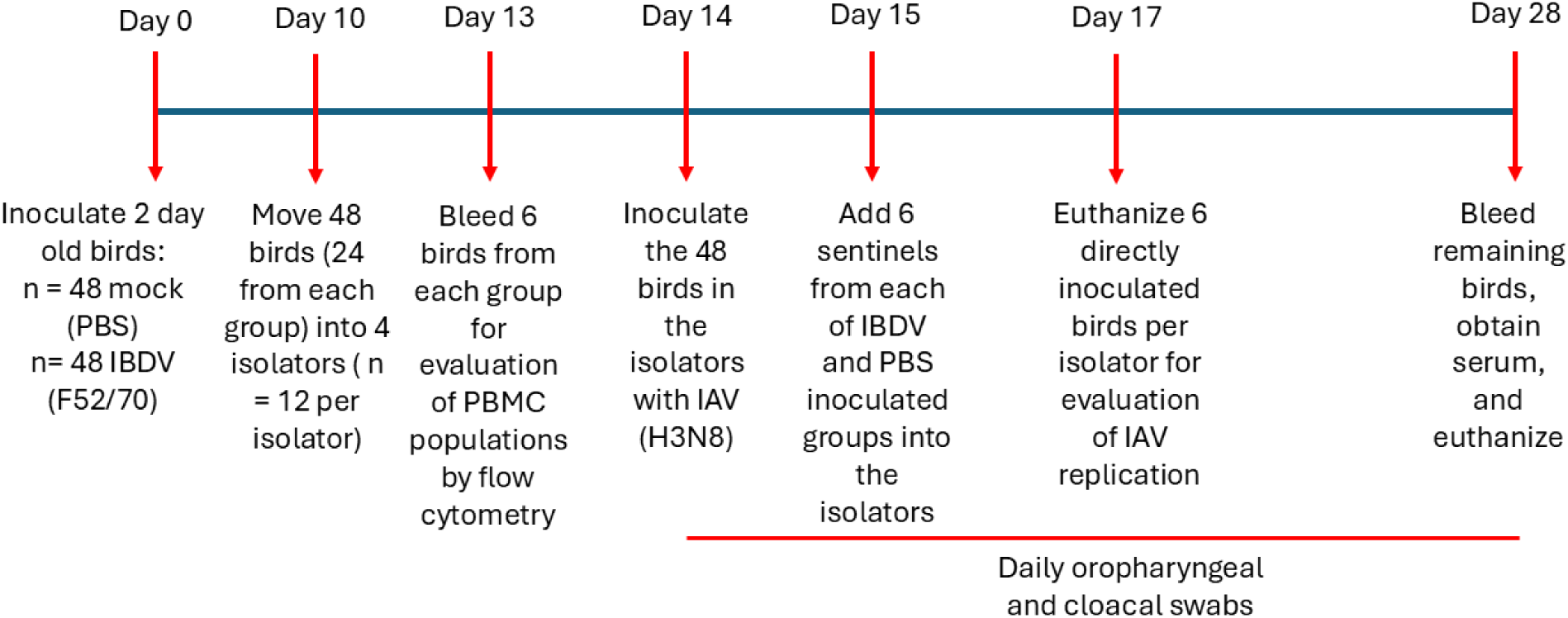
Experimental Design. At two days of age, 96 chickens were randomly allocated to two rooms of 48 birds each. One group was mock-inoculated with PBS, while the other was inoculated with IBDV strain F52/70. At 10 dpi, 24 birds from the mock-inoculated group and 24 birds from the IBDV-inoculated group were divided into 4 isolators, such that isolator 1 and 2 contained 12 mock-inoculated birds each, and isolators 3 and 4 contained 12 IBDV-inoculated birds each, in addition to the 24 mock and 24 IBDV-inoculated birds that remained in the rooms. At 13 dpi, six birds from each group were bled, and PBMCs were obtained to assess the level of immune dysregulation caused by IBDV, by conducting flow cytometry. At 14 dpi, birds in the isolators (n = 48 total) were inoculated with IAV strain A/Mallard/Alberta/156/01 (H3N8). In addition, 12 birds in each room were mock inoculated with PBS, and the remaining 12 in each room served as sentinels. At 15 dpi, six sentinel birds from the mock group were transferred to isolator 1, and six were transferred to isolator 3, while six sentinel birds from the IBDV-inoculated group were transferred to isolator 2, and six were transferred to isolator 4, to make the following 6 groups: Mock/Mock (n = 12) in room 1, IBDV/Mock (n = 12) in room 2, Mock/AIV (n = 12) + mock sentinels (S^mock^) (n = 6) in isolator 1, Mock/AIV (n = 12) + IBDV-infected sentinels (S^IBDV^) (n = 6) in isolator 2, IBDV/AIV (n = 12) + S^mock^ (n = 6) in isolator 3, and IBDV/AIV (n = 12) + S^IBDV^(n = 6) in isolator 4. Oropharyngeal and cloacal swabs were collected from all birds in each isolator from 14 dpi to 28 dpi with IBDV (0-14dpi with IAV) to determine IAV shedding. At 17 dpi with IBDV (3 dpi with IAV), six birds from each group were humanely euthanized and the amount of IAV replicating in tissue samples quantified. The remaining birds were culled at 28 dpi with IBDV (14dpi with IAV) and anti-IAV antibodies in the serum quantified by HI assay.

**Fig 2.**
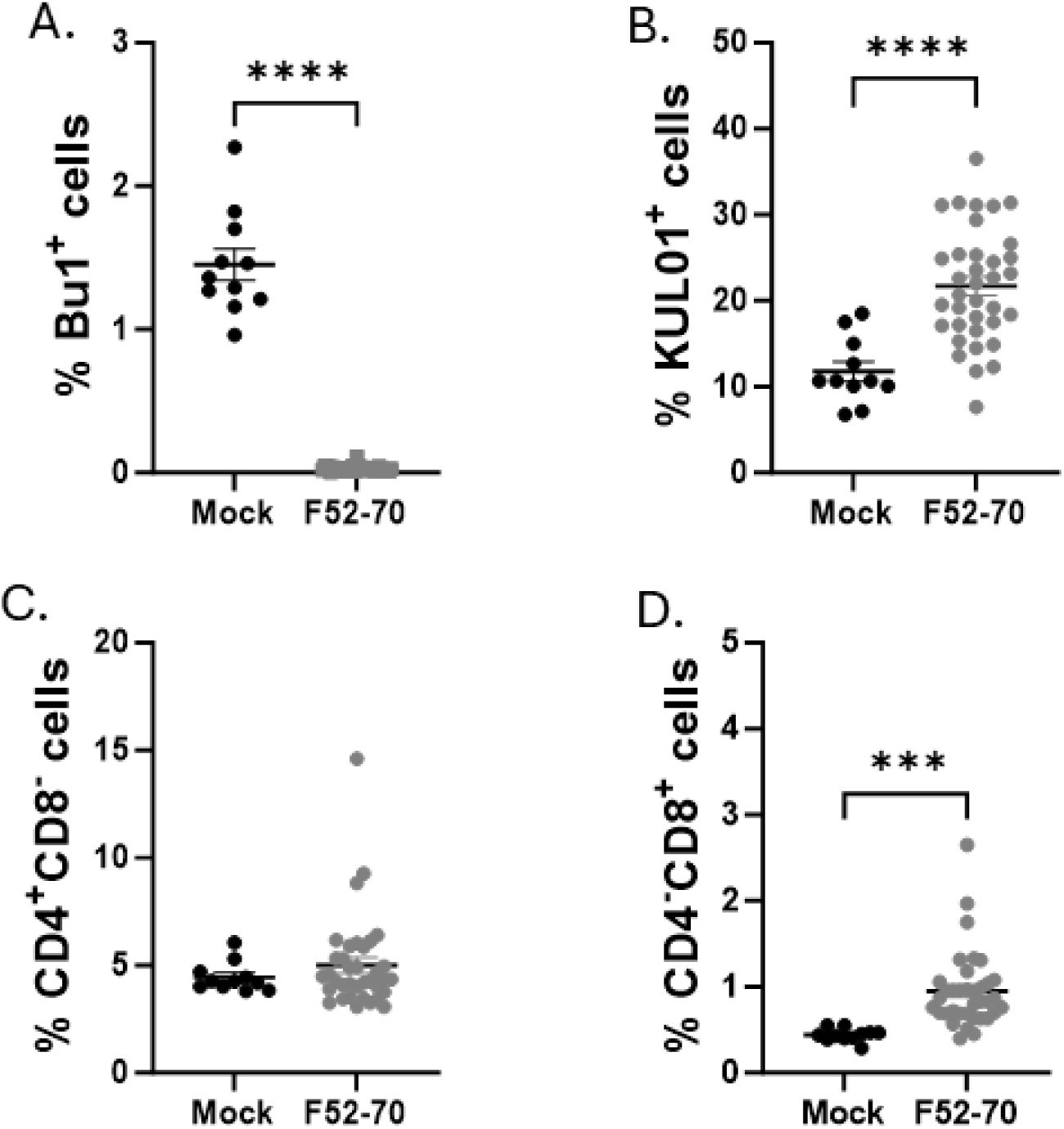
IBDV infection caused immune dysregulation in chickens that were subsequently challenged with the mallard IAV. One day prior to IAV inoculation, blood samples were obtained, and PBMCs were isolated and processed for flow cytometry to quantify the immune cells present. The percentage of Bu1^+^ cells (B cells) (A), KULO1^+^ cells (monocytes/macrophages) (B), CD4^+^CD8^-^ T cells (C), and CD4^-^CD8^+^ T cells (D) were quantified in the samples from the Mock group (black) and IBDV group (grey). The Live/Dead-Fixable Near IR stain was used for dead cell exclusion. The bars represent the mean and standard error of the mean for each population, and the asterisks indicate a statistically significant difference between the averages (*** indicates p ≤ 0.001, and **** indicates p ≤ 0.0001).

### IBDV infection did not significantly alter the clinical progression of the mallard H3N8 IAV in chickens

Infection of the RIR chickens at 2 days of age with IBDV (F52/70) and at 14 days of age with the mallard H3N8 IAV caused mild clinical signs. Neither severe disease, nor mortality was observed in any group, and no bird reached its humane end-point, in keeping with prior studies of IBDV infection of this strain in this age of birds (22), and consistent with the mallard H3N8 IAV being a low pathogenicity avian influenza (LPAI) strain. Birds were weighed daily, and the average daily weight gain was calculated (Fig 3). At 28dpi, there was no significant difference in the average daily weight gain between the Mock/Mock group (17.0 g/day SD +/- 2.6) and the Mock/IAV group (15.3 g/day +/- 1.6), p= 0.27, demonstrating that the IAV infection did not cause a significant reduction in average daily weight gain, consistent with it being a LPAI. In contrast, IBDV infection caused a significantly reduced average daily weight gain in both the IBDV/Mock group (13.5 g/day SD +/- 1.6) (p <0.01) and the IBDV/IAV group (13.8 g/day SD +/- 2.4) (p <0.01) compared to the Mock/Mock group. However, there was no significant difference in the average daily weight gain between Mock/IAV group and the IBDV/IAV group p = 0.20, demonstrating that IBDV infection did not significantly alter the average daily weight gain observed following LPAI virus infection. Taken together, IBDV did not significantly alter the clinical progression of the mallard H3N8 IAV in chickens.

**Fig 3.**
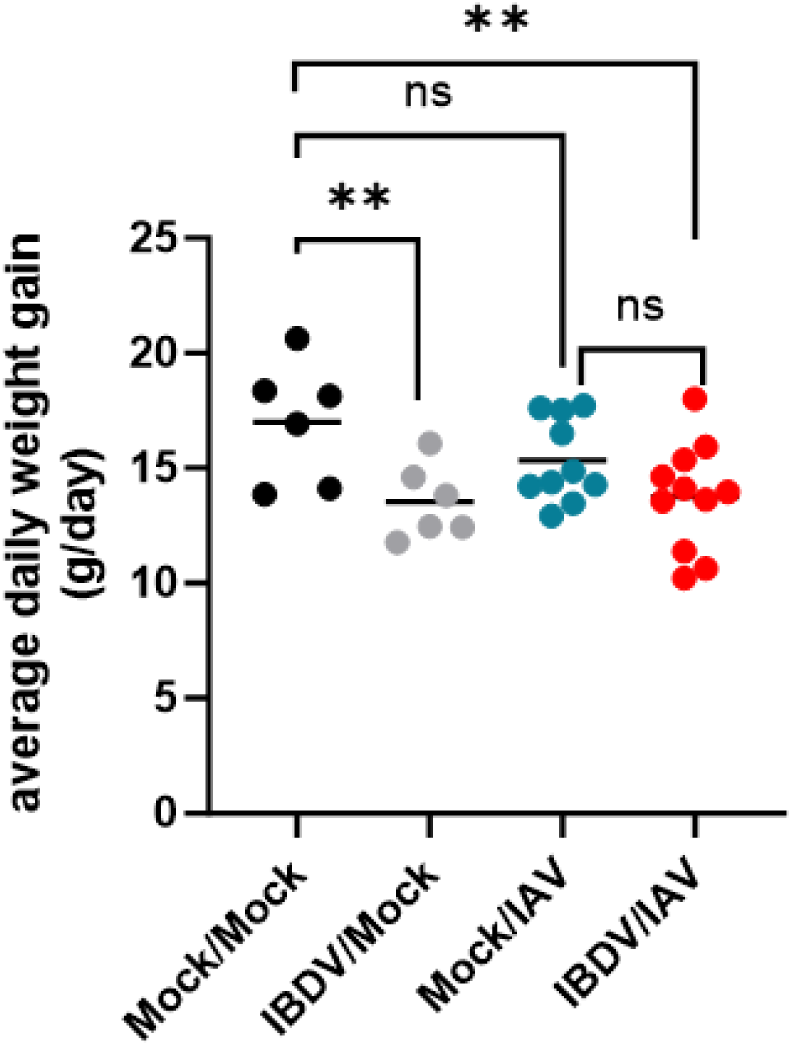
IBDV infection did not significantly alter the average daily weight gain of chickens inoculated with the mallard IAV strain. Birds were weighed daily, and at the final day of the study (28dpi with IBDV), the average daily weight gain was compared between the Mock/Mock group (black), IBDV/Mock group (grey), Mock/IAV group (blue) and IBDV/IAV group (red). The bars represent the mean for each population, and the asterisks indicate a statistically significant difference between the averages (** indicates p ≤ 0.01, ns = not significant).

### IBDV infection increased the average replication of the mallard H3N8 IAV in the trachea of chickens, although this did not reach statistical significance

At 17dpi with IBDV (3dpi with IAV), 12 chickens from the Mock/IAV group and 12 from the IBDV/IAV group were humanely euthanized, and the amount of IAV replicating in the nasal epithelium, trachea, lung, colon, and kidney was quantified by RTqPCR. IBDV infection did not lead to statistically significant changes in IAV replication in the tissue samples, however, the average fold change in the expression of the IAV gene segment M was higher in the trachea of the IBDV/IAV group (average fold change 21.2/mg tissue) compared to the Mock/IAV group (average fold change 0.41/mg tissue), suggesting that IBDV infection could be increasing replication in the trachea (Fig 4).

**Fig 4.**
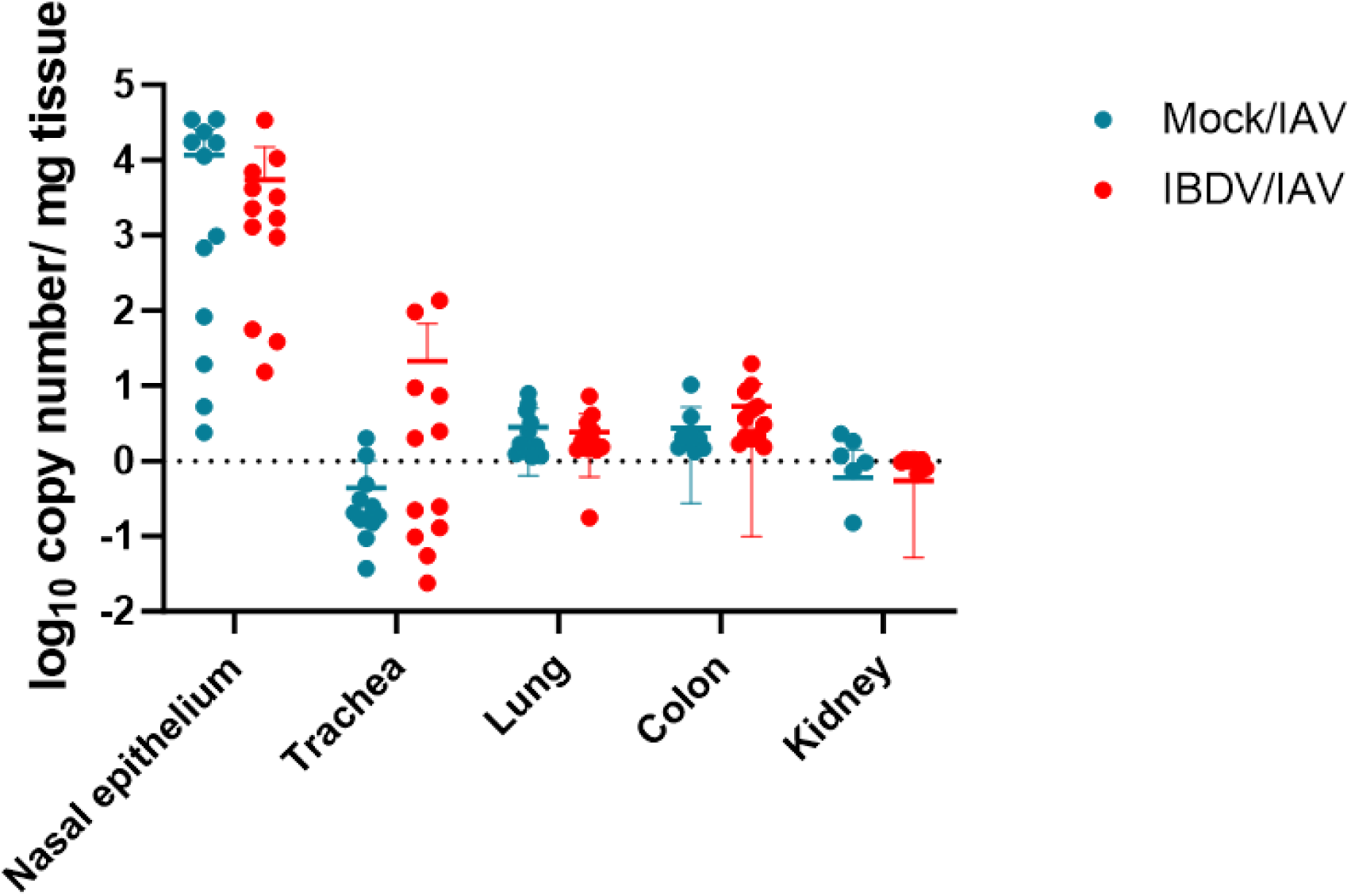
IBDV infection did not cause significant changes in IAV replication in tissues, but did increase the average replication of the mallard LPAI virus in the trachea of chickens. At 17dpi with IBDV (3dpi with IAV), 12 chickens from the Mock/IAV group and 12 from the IBDV/IAV group were humanely euthanized and the amount of IAV replicating in the nasal epithelium, trachea, lung, colon, and kidney was quantified by RTqPCR. The Log_10_ fold change in the expression of the viral M gene segment per sample was quantified on the basis of a standard curve, and expressed as fold change per mg tissue. Each dot represents one sample, Mock/IAV group in blue, and IBDV/IAV group in red. The bars represent the mean and standard deviation for each population.

### IBDV infection prolonged the shedding of the mallard H3N8 IAV from the oropharyngeal cavity of chickens

Oropharyngeal and cloacal swabs were collected from all birds in the Mock/IAV and IBDV/IAV groups from 14-28dpi with IBDV (0-14 dpi with IAV). Swabs were placed in viral transport media, and the amount of infectious IAV in the swab samples quantified by plaque assay on MDCK cells. No infectious IAV was detected in any of the cloacal swab samples, whereas infectious IAV was detected in the oropharyngeal swab samples in over 93% of the chickens at day 1 post infection (Fig 5A), with a mean peak titer of 559 PFU/mL in the Mock/IAV group and 826 PFU/mL in the IBDV/IAV group, but there was no significant difference in the titer between the groups (p = 0.29) (Fig 5B). Chickens with no prior IBDV exposure cleared the IAV within 6 days: infectious IAV could only be detected in one sample at 5dpi (titer = 10 PFU/mL), with none at 6dpi. In contrast, chickens that had been infected with IBDV prior to the mallard LPAI virus (IBDV/IAV), shed the IAV for longer, clearing virus in 7 days: infectious IAV was present in 2 samples at 5dpi (titers = 100 and 120 PFU/mL), one sample at 6dpi (titer = 40 PFU/mL), and none at 7dpi (Fig 5A and B). No infectious IAV was detected in either oropharyngeal or cloacal swab samples from the sentinel birds, and no sentinel birds seroconverted, determined by HI assay, demonstrating that no virus was transmitted from the IAV-inoculated chickens to recipient birds.

**Fig 5.**
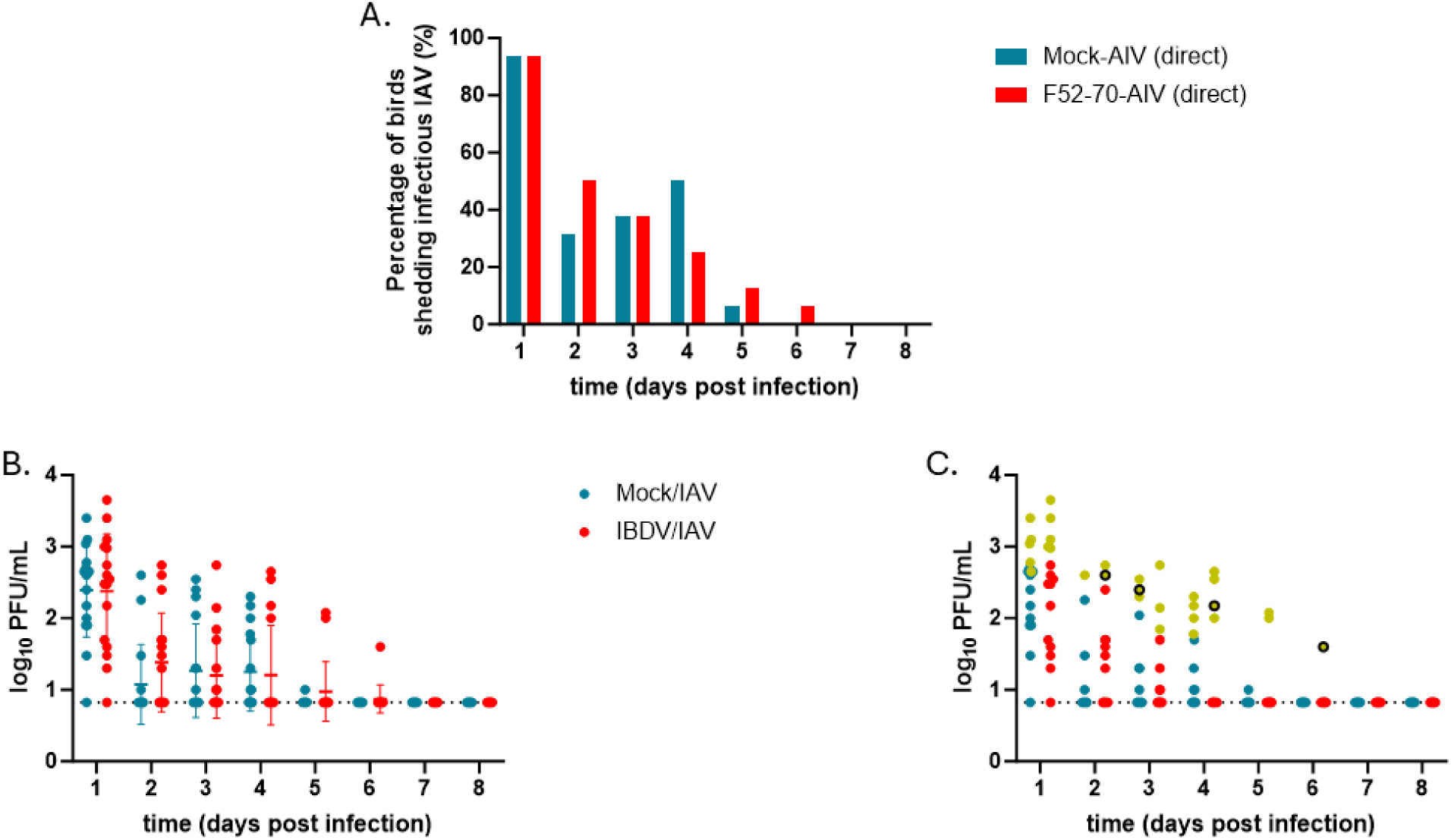
IBDV infection prolonged the shedding of the mallard LPAI virus from the oropharyngeal cavity of some chickens. Oropharyngeal and cloacal swabs were collected from all birds in the Mock/IAV and IBDV/IAV groups from 14-28dpi with IBDV (0-14 dpi with IAV). Swabs were placed in viral transport media, and the titer of infectious IAV in each swab sample quantified by plaque assay on MDCK cells. The percentage of samples that were IAV-positive in at each time point was quantified and plotted, Mock/IAV group in blue and IBDV/IAV group in red (A) and the titer of each sample was expressed as log_10_ plaque forming units (PFU)/mL of swab sample fluid, and plotted (B). Each dot represents one sample, with the Mock/IAV group in blue and the IBDV/IAV group in red. The bars represent the mean and standard deviation for each population. The samples selected for sequencing were highlighted in yellow, and samples where the PCR failed were emphasized with a black border (C).

### IBDV infection increased the mean nucleotide (π) diversity of the mallard LPAI virus population in the chickens, although this did not reach statistical significance

To assess the intra-host evolution of the mallard LPAI H3N8 strain in inoculated chickens, oropharyngeal swab samples that were positive for IAV by plaque assay were selected from nine birds from the Mock/IAV group, and nine birds from the IBDV/IAV group, based on the samples that had the highest titer of IAV from each group. This amounted to 13 swab samples for the Mock/IAV group, and 17 swab samples from the IBDV/IAV group (Fig 5C, yellow dots). However, the RT-PCR reaction failed for one sample in the Mock/IAV group (day 3) and three samples in the IBDV/IAV group (day 2, day 4, and day 6), leaving NGS data from 12 swab samples from 8 birds from the Mock/IAV group (five from day 1, one from day 2, two from day 3, and four from day 4), and 14 swab samples from 8 birds from the IBDV/IAV group (five from day 1, one from day 2, three from day 3, three from day 4, and two from day 5) (Fig 5C and S1 Table).

The fluid from each swab sample was inoculated into embryonated eggs to amplify the virus. The allantoic fluid of the samples, and of the inoculum, was subject to RT-PCR to amplify the IAV genome, which was then sequenced by next generation sequencing (NGS) with an Illumina MiSeq. The sequences of the IAV in the inoculum and the swab samples were then aligned to the reference sequences of the 8 genome segments of the A/Mallard/Alberta/156/01 (H3N8) strain (Accession numbers CY004702.1- CY004709.1), and variants were called from the resulting alignments. The inoculum contained 5 nucleotide changes compared to the reference sequences: PB2 A1220G, HA T1016C, HA T1020A, NA G79A, and NS2 T27C, of which only one, PB2 A1220G, led to an amino acid substitution, PB2 E407G. To assess IAV diversity at the genome level in inoculated birds, we compared the π-diversity between the Mock/IAV and IBDV/IAV groups. IBDV infection increased the mean π-diversity of the mallard LPAI viruses in the chickens, although this did not reach statistical significance (Fig 6), suggesting that IBDV-mediated immune dysregulation had a modest effect on the nucleotide diversity of the population of the mallard strain of IAV in the chickens.

**Fig 6.**
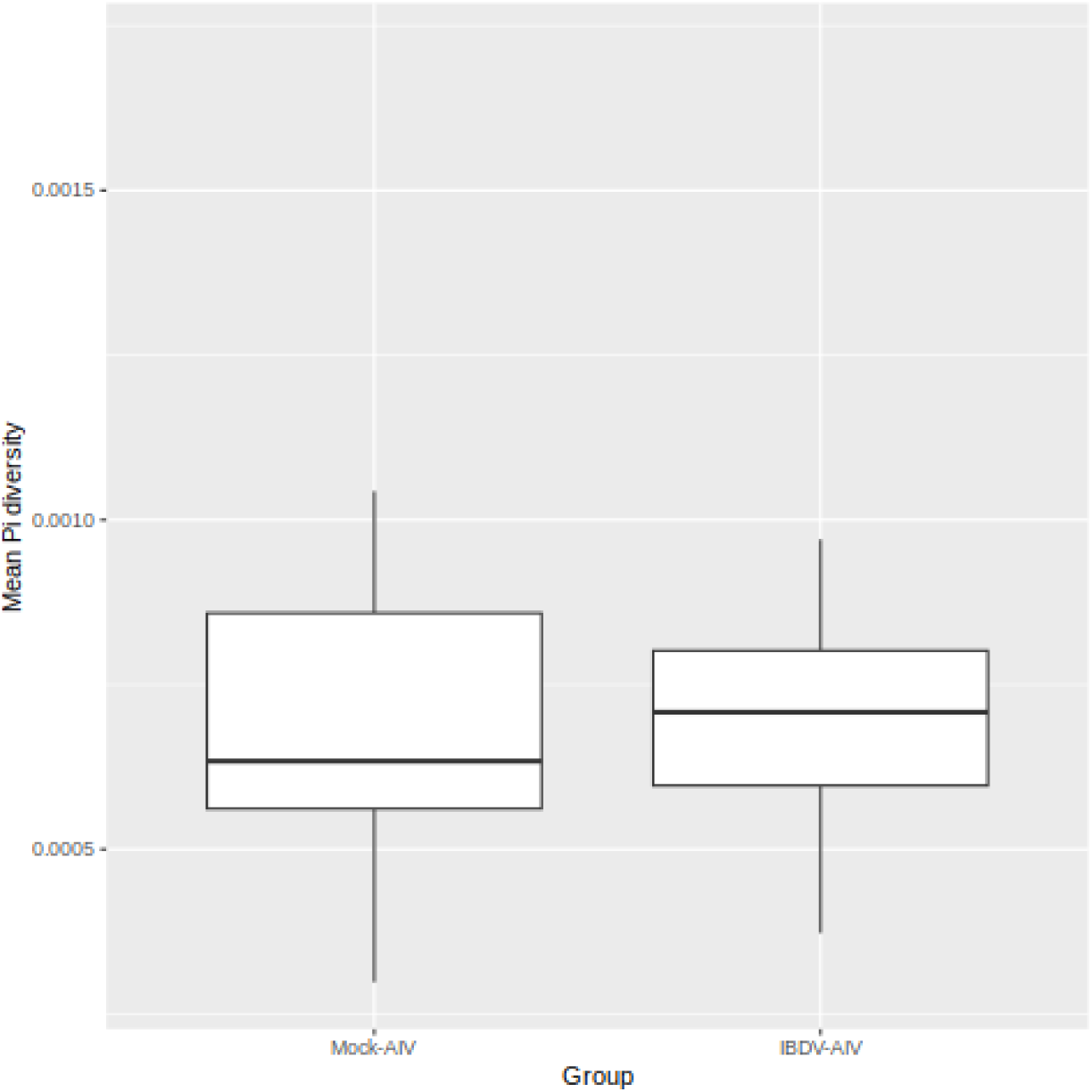
IBDV infection increased the mean nucleotide (π) diversity of the mallard LPAI virus population in the chickens, although this did not reach statistical significance. The fluid from each swab sample selected for sequencing was inoculated into embryonated eggs and the allantoic fluid was harvested 72 hours later, and subject to RT-PCR to amplify the IAV genome, which was then sequenced by NGS with an Illumina MiSeq. The sequences of the IAV in the inoculum and the swab samples were then aligned to the reference sequences of the 8 genome segments of the A/Mallard/Alberta/156/01 (H3N8) strain (Accession numbers CY004702.1-CY004709.1), and variants were called from the resulting alignments. To assess IAV diversity at the genome level in inoculated birds, the π-diversity was calculated and compared between the Mock/IAV and IBDV/IAV groups.

### IBDV infection increased the number of amino acid substitutions detected in the mallard H3N8 IAV population in the chickens

A total of 23 missense variants that led to amino acid substitutions were found in the Mock/IAV samples and 43 amino acid substitutions were detected in the IBDV/IAV samples when all the sequencing reads were considered (S1 Table). When a sequencing depth cut-off of 1,000 sequencing reads was applied to the polymorphism analysis, 13 amino acid substitutions were identified in the Mock/IAV group and 30 were identified in the IBDV/IAV group (Tables 1-3). Prior IBDV infection therefore increased the number of amino acid substitutions detected in the mallard H3N8 population in the chicken hosts.

**Table 1.**
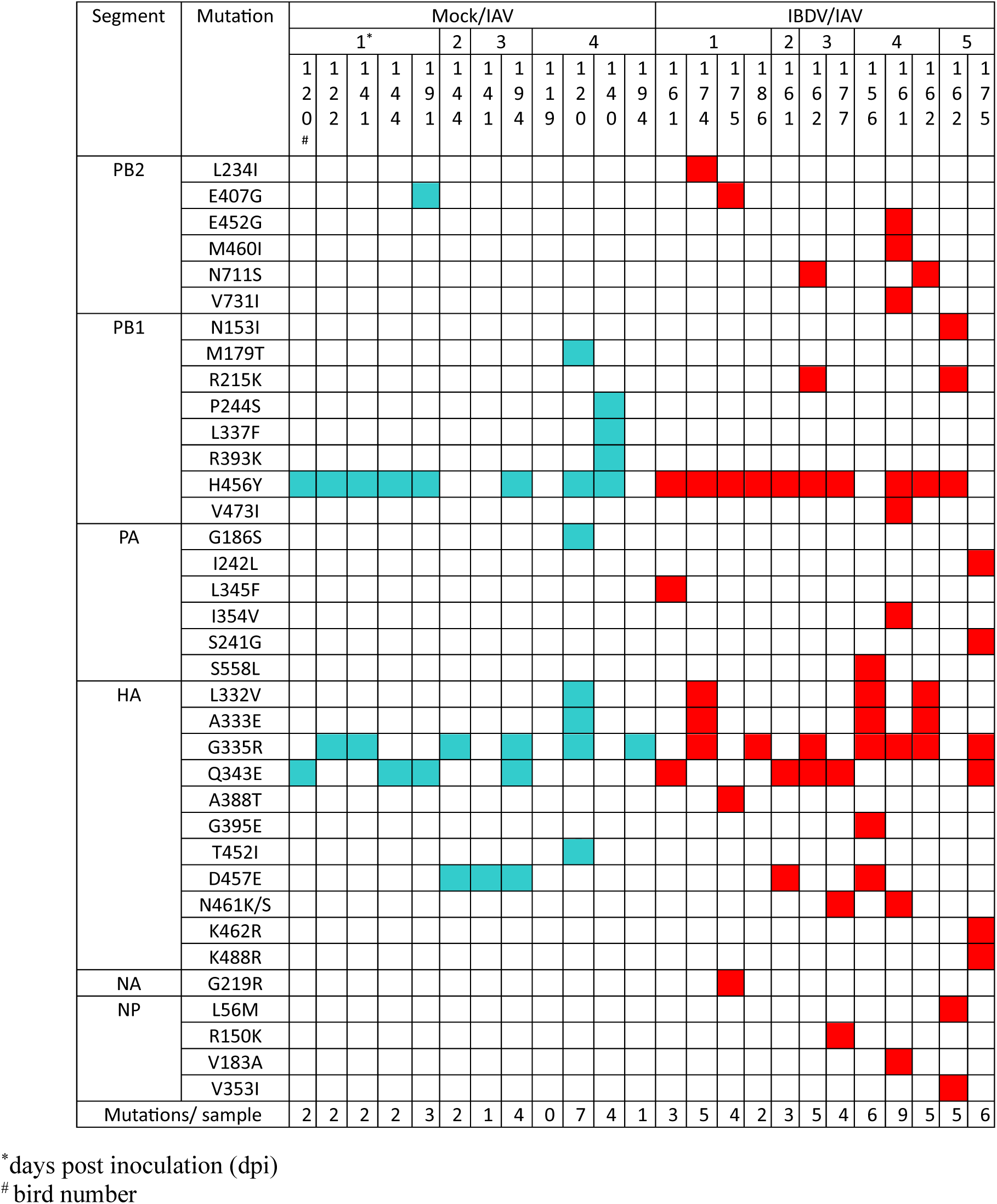
Amino acid substitutions present in the Mock/IAV and IBDV/IAV groups.

### Replication of the mallard H3N8 IAV in chickens was associated with amino acid substitutions in the polymerase complex and HA

Of the 13 amino acid substitutions detected in the Mock/IAV group, the mutations were found in 4 proteins: one in PB2, five in PB1, one in PA, and six in HA (Table 1 blue squares, and Table 2), suggesting that the polymerase complex and HA were associated with adaptation of the mallard virus to the chicken host. Mutations increased as the infection progressed, with four mutations observed 1dpi and ten observed 4dpi (Tables 1 and 2), and 9 of the mutations reached consensus level (defined as a frequency of 0.50 or more) in at least one sample (Table 2). In addition, the majority (8/12 (67%)) of the samples contained multiple amino acid substitutions in both the polymerase complex and the HA, and the average number of mutations per sample was 2.50 (SD +/- 1.83) (Table 1). Out of the 12 samples, PB1 H456Y was found in 8 (67%), HA G335R was found in 6 (50%), HA Q343E was found in 4 (33%), and HA D457E was found in 4 (33%) (Tables 1 and 2). Of these four amino acid substitutions, PB1 H456Y reached consensus level in three of the samples (38%), whereas HA G335R and HA D457E reached consensus level in one sample each (Table 2). Furthermore, of these amino acid substitutions, PB1 H456Y, HA G335R and HA Q343E were found in samples as early as 1 dpi (Tables 1 and 2). AlphaFold 3 modeling revealed that the PB1 H456Y amino acid substitution was exposed on the surface of the polymerase complex and not buried within (Fig 7A), and the HA G335R, Q343E, and HA D457E substitutions were present in the stem region of the HA (Fig 7B and C). None of the HA mutations were predicted to alter N- glycosylation.

**Fig 7.**
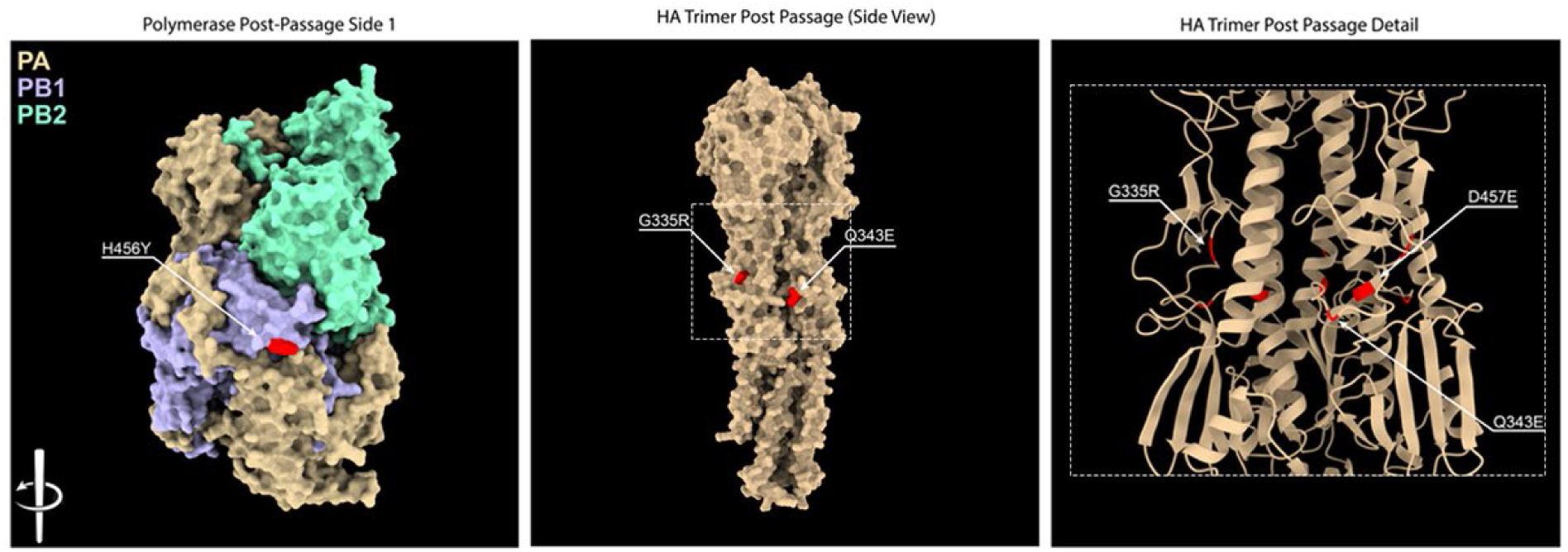
The PB1 H456Y amino acid substitution was exposed on the surface of the polymerase complex, and the HA G335R, Q343E, and HA D457E substitutions were present in the stem region of the HA. The heterohexamer polymerase complex was modeled using the amino acid sequences of the PB2, PB1 and PA from the A/Mallard/Alberta/156/01 (H3N8) strain (Accession numbers: CY004707.1, CY004708.1, and CY004709.1) plus the PB1 H456Y substitution, using AlphaFold3, and a single trimer of PB2, PB1, and PA was selected and rendered with UCSF Chimera X, with the substitution highlighted in red (A). The HA homotrimer was modelled using the amino acid sequence of the HA from the A/Mallard/Alberta/156/01 (H3N8) strain (Accession number: CY004702.1) plus the HA G335R, Q343E, and HA D457E substitutions, using AlphaFold3, and the structure was rendered with UCSF Chimera X, with the substitutions highlighted in red (B). The boxed region of (B) was then depicted as a ribbon diagram, to better emphasize the HA D457E substitution (C).

**Table 2.** Amino acid substitutions present in the Mock/IAV group.

| Gene segment | Amino acid position | Amino acid substitution | 1dpi* | 2 dpi | 3dpi | 4dpi | Total (%) | Variant frequency (dpi) | Read depth |
| --- | --- | --- | --- | --- | --- | --- | --- | --- | --- |
| PB2 | 407 | E -> G | 1 | 0 | 0 | 0 | 1/12 (8%) | 0.24 | 1,798 |
| PB1 | 179 | M -> T | 0 | 0 | 0 | 1 | 1/12 (8%) | 0.81 | 1,642 |
|  | 244 | P -> S | 0 | 0 | 0 | 1 | 1/12 (8%) | 1.00 | 2,724 |
|  | 337 | L -> F | 0 | 0 | 0 | 1 | 1/12 (8%) | 0.32 | 4,064 |
|  | 393 | R -> K | 0 | 0 | 0 | 1 | 1/12 (8%) | 0.75 | 5,440 |
|  | 456 | H -> Y | 5 | 0 | 1 | 2 | 8/12 (67%) | 0.34 (1dpi) | 4,752 |
|  |  |  |  |  |  |  |  | 0.42 (1dpi) | 5,691 |
|  |  |  |  |  |  |  |  | 0.26 (1dpi) | 7,250 |
|  |  |  |  |  |  |  |  | 0.56 (1dpi) | 8,768 |
|  |  |  |  |  |  |  |  | 0.29 (1 dpi) | 4,498 |
|  |  |  |  |  |  |  |  | 0.45 (3 dpi) | 16,430 |
|  |  |  |  |  |  |  |  | 0.74 (4 dpi) | 16,098 |
|  |  |  |  |  |  |  |  | 1.00 (4 dpi) | 6,979 |
| PA | 186 | G -> S | 0 | 0 | 0 | 1 | 1/12 (8%) | 0.21 | 19,262 |
| HA | 332-333 | LA -> VE | 0 | 0 | 0 | 1 | 1/12 (8%) | 0.26 | 1,467 |
|  | 335 | G -> R | 2 | 1 | 1 | 2 | 6/12 (50%) | 0.50 (1dpi) | 1,179 |
|  |  |  |  |  |  |  |  | 0.48 (1dpi) | 1,480 |
|  |  |  |  |  |  |  |  | 0.49 (2dpi) | 1,445 |
|  |  |  |  |  |  |  |  | 0.38 (3dpi) | 1,214 |
|  |  |  |  |  |  |  |  | 0.47 (4dpi) | 1,161 |
|  |  |  |  |  |  |  |  | 0.39 (4dpi) | 2,607 |
|  | 343 | Q -> E | 3 | 0 | 1 | 0 | 4/12 (33%) | 0.33 (1dpi) | 1,355 |
|  |  |  |  |  |  |  |  | 0.25 (1dpi) | 2,326 |
|  |  |  |  |  |  |  |  | 0.20 (1dpi) | 1,751 |
|  |  |  |  |  |  |  |  | 0.33 (3dpi) | 3,712 |
|  | 452 | T -> I | 0 | 0 | 0 | 1 | 1/12 (8%) | 0.77 | 79,546 |
|  | 457 | D -> E | 0 | 1 | 2 | 0 | 3/12 (25%) | 0.43 (2dpi) | 73,709 |
|  |  |  |  |  |  |  |  | 0.54 (3dpi) | 28,252 |
|  |  |  |  |  |  |  |  | 0.22 (3dpi) | 122,054 |

\*days post inoculation (dpi)

### IBDV infection caused changes to the intra-host evolution of the mallard H3N8 virus in the chicken host

Of the 30 amino acid substitutions detected in the IBDV/IAV group, the mutations were found in 6 proteins: six in PB2, four in PB1, five in PA, ten in HA, one in NA, and four in NP (Table 1 red squares, and Table 3). Ten substitutions were detected 1dpi, greater than in the Mock/IBDV group, suggesting that IBDV infection led to a rapid accumulation of substitutions in the IAV. IBDV infection also increased the number of mutations reaching consensus level in at least one sample from 9 in the Mock/IAV group to 18 in the IBDV/IAV group (Table 3), however, as with the Mock/IAV group, the majority of the amino acid substitutions were found sporadically in the samples. The same amino acid substitutions that were detected more frequently in the Mock/IAV samples were also detected in the IBDV/IAV samples, but there were differences in the percentage of the samples in which they were detected: Out of the 14 samples, PB1 H456Y was found in 10 (71%), HA G335R was found in 7 (50%), HA Q343E was found in 5 (36%), and HA D457E was found in 2 (14%). Of these four amino acid substitutions, PB1 H456Y reached consensus level in five of the samples (50%), HA G335R in five of the samples (83%), and D457E in one of the samples (Table 3). Moreover, the frequency of the reads with the HA G335R substitution was significantly higher in the IBDV/IAV group compared to the Mock/IAV group (p<0.05) (Table 3). Furthermore, of these amino acid substitutions, PB1 H456Y, HA G335R and HA Q343E were also found in samples as early as 1 dpi (Tables 1 and 3). Finally, IBDV infection also increased the average number of mutations per sample from 2.50 (SD +/- 1.83) in the Mock/IAV group to 4.75 (SD +/- 1.81) in the IBDV/IAV group (p < 0.01) (Table 1 and Fig 8).

**Fig 8.**
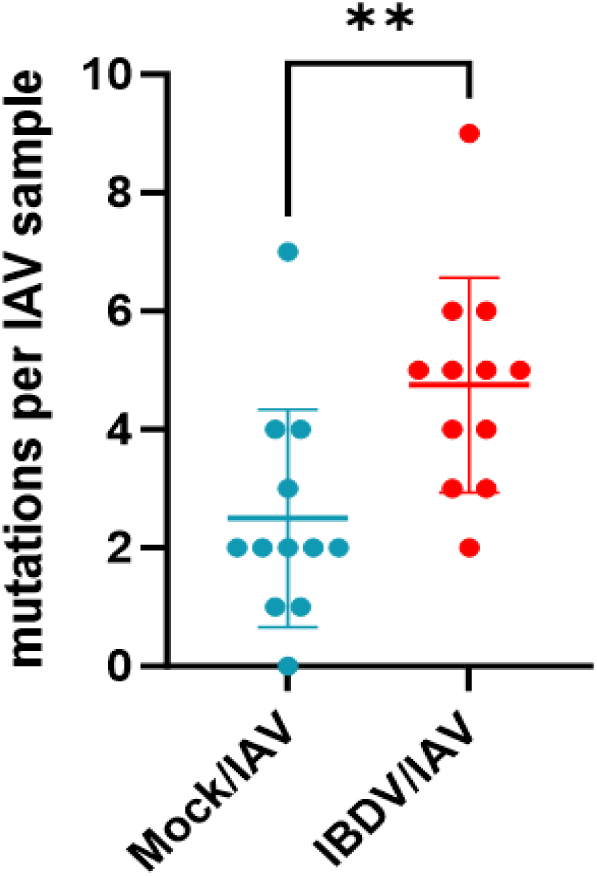
IBDV infection increased the average number of mutations per IAV sample. The number of IAV amino acid substitutions detected in each sample was calculated and plotted for the Mock/IAV and IBDV/IAV groups. Each dot represents one sample. The bars represent the mean and standard deviation for each group and the asterisks indicate a statistically significant difference between the averages (** indicates p ≤ 0.01).

**Table 3.**
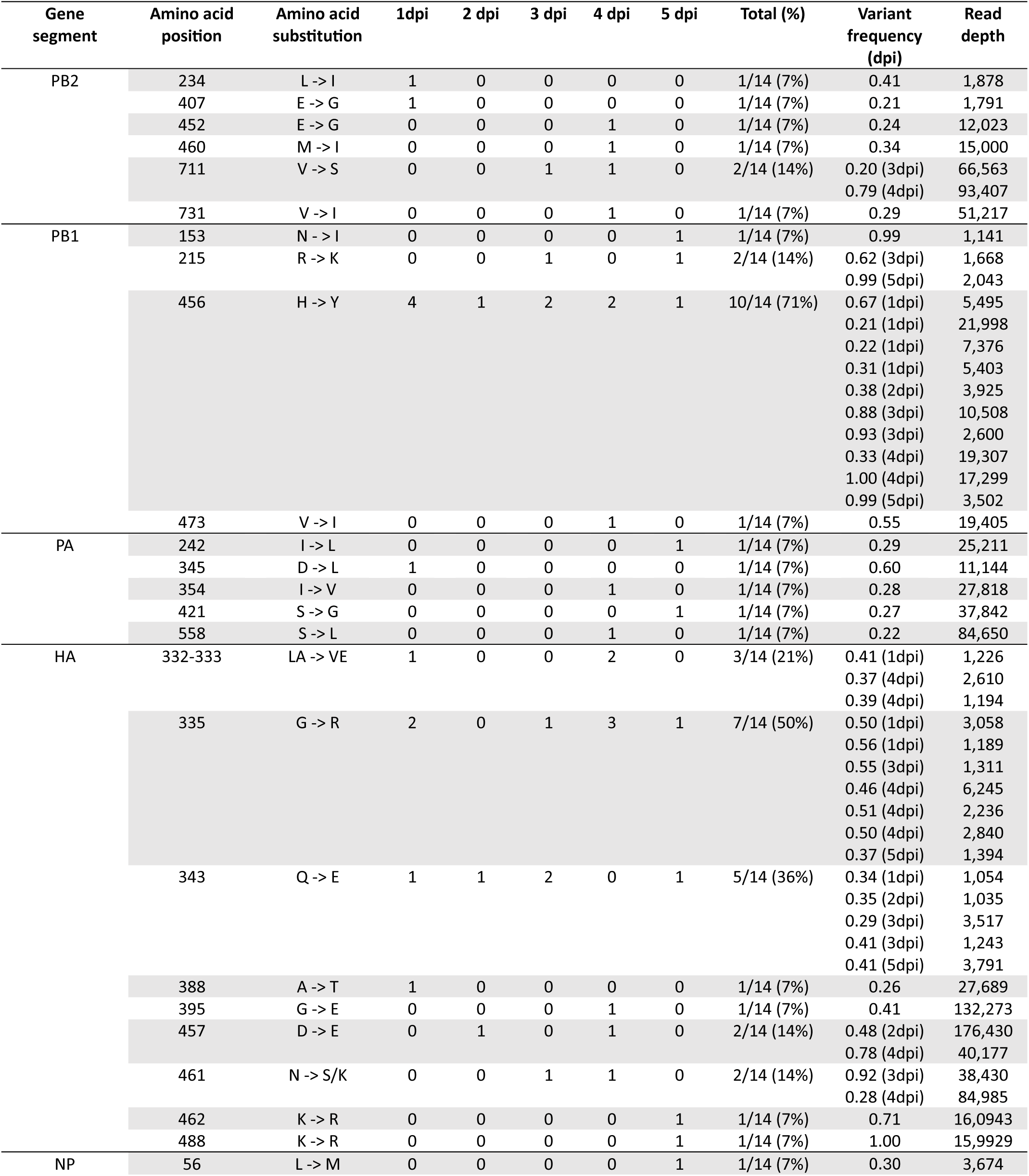
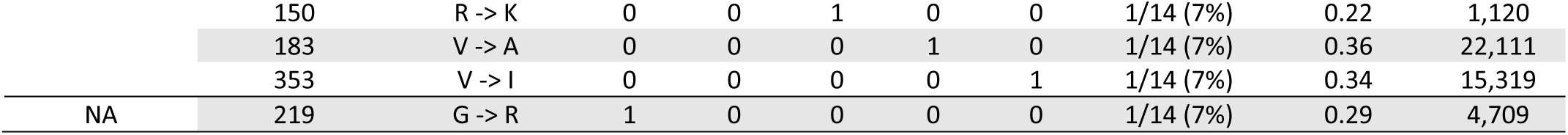
Amino acid substitutions present in the IBDV/IAV group.

Taken together, these data indicate that adaptation of the mallard H3N8 virus to the chicken host was associated with amino acid substitutions in the polymerase complex and HA, and that IBDV-mediated immune dysregulation caused changes to the intra-host evolution of the IAV, significantly increasing the number of mutations per IAV sample, increasing the total number of amino acid mutations in the IAV population, increasing the number of viral proteins in which mutations were found, and increasing the number of mutations reaching consensus level.

## Discussion

In this study, we inoculated chickens with an IAV strain isolated from a mallard (A/Mallard/Alberta/156/01 (H3N8)) (23) and compared the replication and shedding of the IAV between groups of chickens that were either immunosuppressed by an IBDV infection, or immunocompetent controls. We selected this strain of IAV as we wanted to use a strain that had been isolated from aquatic waterfowl and not passaged in chickens that we could work with in an ABSL2 environment. Moreover, it has previously been reported that LPAI strains sampled from Anseriformes in Alberta were representative of AIVs circulating in North American Anseriformes (24). Our experimental design involved the use of SPF chickens of the RIR line that were immunosuppressed by intranasal inoculation with 10^5^ TCID_50_/bird of IBDV strain F52/70 at 2 days of age. We selected this experimental design as this IBDV strain, dose, and route of inoculation in this breed and age of chickens has previously been shown to induce a lasting immunosuppression (22). Moreover, we wanted to model a situation that might be encountered in a backyard setting where chicks might have been obtained from a flock that had not been vaccinated against IBDV, and therefore may be immunologically naïve and susceptible to IBDV infection at an early age, given that IBDV is endemic globally. We then challenged the birds with the mallard IAV strain at 14 days post-IBDV inoculation. This time frame was selected as we wanted the birds to have recovered clinically from the IBDV infection prior to IAV challenge to avoid confounding the results of the IAV challenge. Moreover, given the sporadic nature of IAV spillover from wild birds into domestic poultry, we reasoned that chickens may be older when they become exposed to IAV, compared to when they might be exposed to IBDV in the field.

To confirm that IBDV caused immune dysregulation at the time of IAV inoculation, we performed flow cytometry. IBDV infection significantly reduced the percentage of B cells in the peripheral blood compared to mock-inoculated birds, even at 13dpi. This approach enabled us to evaluate the immune status of the same birds that were subsequently inoculated with IAV at 14dpi, rather than relying on bursal pathology on a sample of the birds as a marker of immunosuppression. These data are consistent with previous reports that demonstrated quantifying peripheral blood B cells is a good correlate of IBDV-induced immunosuppression (25). Moreover, it has previously been demonstrated that infection with this IBDV strain in this age of birds causes a reduction in B cell number in the peripheral blood (22). Interestingly, the percentage of monocytes/macrophages, CD8+, and CD4+CD8+ T cells was increased in the peripheral blood at 13dpi. This demonstrates that the birds had immune dysregulation at the time of IAV challenge, rather than a suppression of all immune cell populations.

Next, we quantified daily weight gain and daily IAV shedding in oropharyngeal and cloacal swabs for 14 days following IAV infection. Additionally, at 3 days post- IAV infection, we quantified the amount of virus replicating in different organs. Moreover, we added immunocompetent or immunosuppressed sentinel birds to the IAV inoculated groups to monitor if any IAV transmission occurred. We found that the mallard H3N8 virus stain did not cause severe disease in inoculated chickens, and did not cause any significant change in weight compared to the mock-inoculated control group, consistent with it being a low pathogenicity virus. The IAV replicated in the upper respiratory tract of the chickens by RTqPCR, and we determined that infectious IAV was shed from the oropharyngeal cavity by plaque assay, but we did not detect any IAV positive cloacal swab samples, indicating that the virus was not shed from the cloaca of inoculated chickens. This is consistent with previous reports that revealed LPAI viruses are rarely shed from the cloaca (26). We also did not detect transmission of the IAV to the sentinel chickens, as no virus was detected in the oropharyngeal or cloacal swabs obtained from the sentinel birds by plaque assay, and none of the sentinel birds had antibodies against IAV by HI assay. These data are consistent with previous reports that avian influenza virus strains transmit inefficiently between chickens, even if they transmit efficiently between ducks (9, 27–29), although the published studies were with H5 and H2 subtypes. In contrast, studies with H7 viruses have yielded mixed results: In one study, two turkey viruses, the LPAI strain A/turkey/Indiana/16-001571-6/2016 (H7N8) and the high pathogenicity avian influenza (HPAI) strain A/turkey/Indiana/16-001403-1/2016 (H7N8) failed to transmit between chickens (30), but in another study, the LPAI strain A/duck/Alabama/2017 (H7N9) did show transmission between chickens (31), demonstrating that the ability of avian influenza viruses to spread between chickens is strain-dependent.

Prior infection of chickens with IBDV prolonged the shedding of the mallard LPAI H3N8 virus challenge from the oropharyngeal cavity from 5 days to 6 days in some chickens and increased the average fold change of IAV replication at 3dpi in the trachea, although this did not reach statistical significance. These data are consistent with a previous report by Khalil *et al.* (2023) that revealed a live IBDV vaccine (strain 228E) given at 17days of age increased the replication of an influenza A/chicken/Egypt/FAO-S33/2021 (H9N2) virus challenge given 5 days later in tracheal swab samples, as determined by RTqPCR (18). In their study, data were expressed as Ct values, and birds in the IBDV/IAV group had lower IAV Ct values at 4 and 8 dpi (29.48 +/-0.73 at 4dpi and 31.73 +/- 0.48 at 8dpi) than birds in the Mock/IAV group (31.57 +/- 0.15 at 4 dpi and 35.5 +/- 0.75 at 8dpi), indicating more IAV shedding (18). Moreover, our data are in agreement with a report by Hashemzade *et al.* (2019) who found that, in turkeys, IBDV inoculation increased the replication of IAV A/chicken/Iran/688/1999 (H9N2) in the trachea from 2.12 log_10_ +/-0.13 in the Mock/IAV group to 4.08 log_10_ fold change +/-0.15 in the IBDV/IAV group at 3dpi. The same trend also occurred in the lung, with IAV replicating to 1.95 log_10_ +/-0.14 in the Mock/IAV group, which increased to 3.84 log_10_ +/-0.22 in the IBDV/IAV group (20). However, our data, and that of Khalil and Hashemzade *et al.* are in opposition to the conclusions made by Ranjbar *et al.* (2019), who reported that very virulent (vv) IBDV infection suppressed IAV replication in the trachea of chickens, which the authors suggested was possibly due to cytokine responses induced by the IBDV interfering with the IAV replication (19). The reason for the differing outcomes remains unknown, although it is important to note that the observed suppression in IAV replication occurred at 3 and 6dpi, whereas at 1dpi, the opposite was observed, and IBDV infection enhanced IAV replication from approximately 20,000 copies of the IAV genome in the trachea of the Mock/IAV group to approximately 120,000 copies of the IAV genome in the trachea of the IBDV/AIV group (19), suggesting that rather than IBDV suppressing IAV replication, IBDV altered the kinetics of IAV shedding, with more IAV shed earlier, and less shed later. Furthermore, vv strains are known to cause severe clinical signs and mortality in infected birds (32), which could have impacted upon the outcome of an IAV challenge. In contrast, we aimed to model the effect of a subclinical immunosuppression in birds that recovered from an IBDV infection and had no clinical signs at the time of IAV challenge, but had proven immune dysregulation. Importantly, the previously documented studies that evaluated the potential for IBDV to exacerbate IAV replication and/or shedding did so using chicken strains of IAV. Therefore, our study is the first to evaluate the effect of IBDV in chickens on the replication and shedding of a challenge with an aquatic waterfowl strain of IAV.

The mechanism of how IBDV increases the replication of IAV in the trachea, or prolongs the shedding of IAV from the oropharyngeal cavity, is unclear. The preferred tropism for IBDV are B lymphocytes, the majority of which reside in the BF, and infection leads to a loss of humoral immunity (33). It is, therefore, possible that IAV-antigen specific humoral responses fail to develop in the IBDV/IAV group compared to the Mock/IAV group, which therefore fail to contain the spread of the virus within the trachea. However, IBDV infection increased the replication of IAV in the trachea at 3dpi in our study, and in the other reported studies IAV replication was elevated in the trachea at 1dpi, 3dpi, or 4dpi due to IBDV (18, 20). Moreover, our study showed IAV shedding was prolonged from 5 to 6 dpi. These time frames are too soon for a loss of IAV antigen-specific humoral responses to be the reason for the difference in IAV replication, especially given that the birds are SPF and have no prior exposure to IAV. Therefore, we hypothesize that IBDV alters the mucosal environment in the respiratory tract in such a way that exacerbates IAV replication or shedding, for example by altering innate immunity or the microbiome, but the mechanism underpinning this remains to be determined.

We hypothesize that the elevated average fold change of IAV in the trachea, and the prolonged shedding of infectious IAV from the oropharyngeal cavity of some chickens could potentially increase the amount of IAV shed into the environment. Although the changes we observed in our study are modest, when scaled up to a flock level, we hypothesize that these modest changes could add up to be more impactful. Importantly, the mallard LPAI H3N8 virus did not transmit to sentinel chickens, suggesting that IBDV-mediated immunosuppression does not increase the risk of IAV between chickens. However, the risk of transmission to other animals, as well as humans remains unknown.

Finally, we sequenced the IAV from the inoculated birds, to determine if there was any intra-host evolution. Replication of the mallard LPAI H3N8 virus in the chicken host was associated with amino acid substitutions in the polymerase-encoding gene segment PB1 (H456Y), and the HA gene segment (G335R, Q343E, and D457E), which were present in 33-67% of the samples in the Mock/IAV group and 14-71% of the samples in the IBDV/IAV group. The presence of the same amino acid substitution across multiple samples from multiple animals is a strong indicator of a selective process (34). Moreover, the amino acid substitutions arose without serial passage in the chickens and could be found as early as 1 dpi (Tables 1-3), demonstrating they arose rapidly. The adaptation of IAV strains from wild birds into poultry is known to be associated with amino acid substitutions in the polymerase complex and the HA (26, 31, 35), and our observations are consistent with these reports. For example, Chrzastek *et al.* infected chickens with a LPAI influenza A/duck/Alabama/2017 H7N9 strain and discovered 45 amino acid substitutions at 4 and 6 dpi, 16 in the PB2 segment, 7 in PB1, 18 in PA, 2 in NS, and 2 in NP (31), demonstrating that the polymerase complex was particularly important in host adaptation. Similarly, Leyson et al inoculated chickens with H7N3 viruses isolated from turkeys that had become infected from wild birds and found that replication in chickens was associated with consensus level amino acid substitutions in the proteins involved in viral replication: 1 in PB1, 1 in PA, 1 in NP, and 2 in NS (29). Moreover, Youk *et al*. 2021 compared the sequences of influenza A/Northern Pintail/Washington/40964/2014 (a duck origin H5N2) and A/Chicken/Iowa/13388/2015 (a chicken origin H5N2), which revealed the viruses differed by 20 amino acids. The presence of these amino acids was then evaluated in all H5N2 viruses isolated in the USA and the mutations that were associated with adaptation to chickens were identified, which were in PB1, NP, HA, and NA (35).

A protein (p)BLAST search of the amino acid sequence of the PB1 gene (Accession number CY004708.1) revealed that out of 100 PB1 sequences, 95 had an H at position 456 and 5 had a Y, indicating some variability at this site. However, the PB1 sequences that had an Y at position 456 were found in wild birds (teal, mallards, and a “wild waterbird” (24)), suggesting that this amino acid substitution was not adaptive for chickens specifically. Interestingly, the PB1 H456Y mutation has previously been reported to arise in the brain of mice inoculated with the influenza A/Hong Kong/156/97 (HK156) strain of H5N1(36), where it was associated with neurotropic adaptation and virulence. Moreover, in 2024, a human patient presenting with symptoms of hemolytic uremic syndrome and a hemorrhagic pneumonia harbored a large population of H1N1 pdm09 viruses with a PB1 H456H/Y substitution (37). Given that the PB1 H456Y mutation is predicted to be exposed on the surface the polymerase complex (Fig 7A), it is tempting to speculate that it could interact with cellular proteins, however, given that it has been found in a diverse range of species of both birds and mammals, it remains to be determined whether it influences host range restriction. Moreover, given that the PB1 H456Y mutation has been associated with disease in mice and humans, it is possible that the mutation influences IAV virulence. However, it was beyond the scope of the present study to evaluate the pathogenicity of the viruses in a mammalian model.

A pBLAST search of the H3 sequence (Accession CY004702.1) revealed that out of 100 sequences, all had a G at position 335 a Q at position 343, and a D at position 457. Outside of the H3 subtype, the equivalent amino acid to position 335 was a G in all subtypes (38), whereas the equivalent amino acid to position 343 was a Q in subtypes H 1,3, 12, 14, and an E in H 2,5, and 8 (38). The equivalent amino acid to position 457 was a D in all subtypes. Given the high frequency of the G335R mutation in our study, it is interesting that a G at this position appears to be conserved across the different HA subtypes. The functional implications of the HA amino acid substitutions remain undetermined. The HA amino acid substitutions were present in the stem region of the molecule, and they were not predicted to alter glycosylation. It is possible that they might alter the ability of the HA to fuse with the endosome during cellular entry, but this hypothesis remains to be tested and an evaluation of whether these mutations altered pH stability, receptor binding, or immune evasion potential was beyond the scope of the current study.

This study is the first to evaluate how IBDV infection affects the intra-host evolution of an IAV challenge. Prior infection with IBDV increased the average nucleotide (π) diversity of the mallard IAV sequences in the chicken samples, but this did not reach statistical significance (Fig 6). IBDV infection also increased the number of amino acid substitutions detected in the IAV population from 13 to 30, and significantly increased the number of mutations per IAV sample from 2.50 (SD +/- 1.83) in the Mock/IAV group to 4.75 (SD +/- 1.81) in the IBDV/IAV group (p < 0.01). IBDV infection also increased the number of IAV proteins in which mutations were found, increased the number of mutations observed in the IAV population at 1dpi and increased the number of mutations reaching consensus level. The reason for these observations remains unknown, but we hypothesize that this could reflect a relaxed selection pressure on the IAV population, allowing it to accumulate more mutations than in an immunocompetent host.

Whether this is due to a loss of humoral immunity or altered innate immunity or microbiome remains to be determined. Interestingly, IBDV-immunosuppressed chickens still shed IAVs containing the PB1 H456Y, HA G335R, HA Q343E, and HA D457E mutations, although the percentage of samples in which the amino acid substitutions were found was altered: PB1 H456Y increased from 8/12 (67%) to 10/14 (71%) of the samples, HA G335R remained the same at 6/12 (50%) and 7/14 (50%) of the samples, HA Q343E increased from 4/12 (33%) to 5/14 (36%) of the samples, and HA D457E decreased from 4/12 (33%) to 2/14 (14%) of the samples. Taken together, these data demonstrate that IBDV infection caused some changes to the intra host evolution of the IAV. We hypothesize that this, together with the prolonged shedding, could potentially lead to the accumulation of a greater diversity of IAVs in the environment when scaled up to a flock level.

Our study is not without limitations. First, we modelled what may happen in backyard flocks, where chicks may hatch with no preexisting immunity to IBDV, become infected with a field strain of IBDV soon after hatch, and then become infected with IAV later. However, commercial flocks are typically vaccinated against IBDV. Vaccinated flocks can still become infected with field strains subclinicaly, which can still lead to immunosuppression, however modeling the effect of IBDV infection in vaccinated birds on the replication, shedding, and intra-host evolution of IAV was beyond the scope of the present study. In addition, we inoculated birds with a classical strain of IBDV (F52/70) that belonged to genogroup A1B1 as we previously demonstrated that this strain of virus caused a lasting immunosuppression in our birds. A1B1 strains circulate in many countries, so our data have real-world significance, however, reassortant strains belonging to genogroup A3B1 are now common in Europe (39, 40) and variant strains belonging to genogroup A2B1 and A2dB1 are common in the US and China, respectively (41–43). Therefore, it would be beneficial in the future to evaluate the effect of different strains of IBDV on the replication, shedding, and intra-host evolution of IAV. Additionally, we modelled the effect of IBDV on a LPAI virus challenge (strain H3N8). Given the ongoing outbreak of AIV subtype H5N1 (clade 2.3.4.4b) since 2022, and the frequent cross-species spillover events that have been observed from wild birds to poultry, it would be beneficial to evaluate how co- morbidities in poultry affect the replication, shedding, and intra-host evolution of a HPAI challenge. Additionally, it would be interesting to evaluate how IBDV immune dysregulation influences the intra-host evolution of LPAI viruses of the H7 and H5 subtypes, which can become HPAI in poultry.

In summary, the adaptation of the mallard LPAI virus to the chicken host was associated with amino acid substitutions in the polymerase and HA proteins. IBDV-mediated immune dysregulation in chickens prolonged the shedding of the IAV challenge in some birds and caused changes to the intra-host evolution of the IAV, significantly increasing the number of amino acid substitutions per IAV sample. We conclude that just as in people, the outcome of influenza infection in birds can be dictated by their comorbidities, and controlling the spread of wild aquatic waterfowl strains of influenza in poultry should involve a holistic approach, including the control of immunosuppressive diseases that could exacerbate their spread, in addition to enhanced biosecurity.

## Materials and Methods

### Cells and Viruses

The immortalized chicken B cell line, DT40 (44), was cultured in Roswell Park Memorial Institute (RPMI) media supplemented with sodium bicarbonate and L-glutamine (Sigma-Aldrich, Merck), 10% heat-inactivated (hi) fetal bovine serum (FBS) (Life Science Production), 1% tryptose phosphate broth (Sigma-Aldrich, Merck), 100 mM sodium pyruvate (Gibco), and 50 mM beta-mercaptoethanol (B-ME) (Gibco). This was named “complete RPMI media”. The maintenance environment was comprised of 37°C and 5% CO_2_. Madin-Darby Canine Kidney cells (MDCK; ATCC CCL-34) were cultured in Dulbecco’s modified Eagle’s medium (DMEM) supplemented with 10% hi FBS and 1% penicillin/streptomycin (Gibco). The classical (c)-strain of IBDV, F-52/70 (genogroup A1B1) (45), was generously provided by Dr. Nicolas Eterradossi (46), the titer of which was quantified by end-point serial dilution in DT40 cells, as previously described (32). IBDV titers were expressed as tissue-culture infectious dose-50/mL (TCID_50_/mL) and calculated according to the method described by Reed and Muench (47). The low pathogenicity avian influenza (LPAI) challenge virus, A/Mallard/Alberta/156/01 (H3N8) (23), was provided by Dr Holly Shelton (The Pirbright Institute) and grown in embryonated hens eggs, the titer of which was quantified by end point serial dilution in MDCK cells, and expressed as plaque forming units/mL (PFU/mL), as previously described (48). The limit of virus detection in the plaque assays was 6.67 PFU/mL.

### Animals

Specific pathogen-free (SPF) chickens of mixed sex belonging to the Rhode Island Red (RIR) breed were sourced from the National Avian Research Facility (NARF) at the Roslin Institute, University of Edinburgh. Birds were shipped at 1 day of age from the Roslin Institute to The Pirbright Institute and housed in either raised floor pens, or BioFlex® B50 Rigid Body Poultry isolators (Bell Isolation Systems, Livingston, UK), maintained at negative pressure, as described below. Chickens were provided with *ad libitum* feed following the manufacturer’s guidelines for their respective age. All animal procedures conformed to the United Kingdom Animal (Scientific Procedures) Act (ASPA) 1986 under Home Office Establishment, Personal and Project licenses, following approval of the internal Animal Welfare and Ethic Review Board (AWERB) at The Pirbright Institute.

### Animal Experiment

At two days of age, 96 chickens were randomly allocated to two groups of 48 birds each (n=48) (Fig 1) and accommodated in separate rooms in raised-floor pens. One group was intranasally inoculated with phosphate-buffered saline (PBS) as a mock control, while the other group received an inoculum of 10^5^ TCID_50_/bird of the IBDV strain F52/70, administered 50μL per nares. At 10 days post-infection (dpi), 24 birds from the mock-inoculated group and 24 birds from the IBDV-inoculated group were divided into 4 self-contained isolators maintained at negative pressure, such that isolator 1 and 2 contained 12 mock-inoculated birds each, and isolators 3 and 4 contained 12 IBDV-inoculated birds each, in addition to the 24 mock-inoculated and 24 IBDV-inoculated birds that remained in the rooms. At 13 dpi, six birds from each group (representing 2 rooms and 4 isolators (n = 36 total) were bled, and peripheral blood mononuclear cells (PBMCs) were obtained to assess the level of immune dysregulation caused by IBDV, by conducting flow cytometry (described below). At 14 dpi, birds in the isolators (n = 48 total) were inoculated with 10^5^ Plaque Forming Units (PFU)/bird of IAV strain A/Mallard/Alberta/156/01 (H3N8), 50μL per nares. In addition, 12 birds in each room were mock inoculated with PBS, and the remaining 12 in each room served as sentinels. At 15 dpi, six sentinel birds from the mock group were transferred to isolator 1, and six were transferred to isolator 3, while six sentinel birds from the IBDV-inoculated group were transferred to isolator 2, and six were transferred to isolator 4, to make the following 6 groups: Mock/Mock (n = 12) in floor pen 1, IBDV/Mock (n = 12) in floor pen 2, Mock/AIV (n = 12) + mock sentinels (S^mock^) (n = 6) in isolator 1, Mock/AIV (n = 12) + IBDV-infected sentinels (S^IBDV^) (n = 6) in isolator 2, IBDV/AIV (n = 12) + S^mock^ (n = 6) in isolator 3, and IBDV/AIV (n = 12) + S^IBDV^(n = 6) in isolator 4. Oropharyngeal and cloacal swabs were collected from all birds in each isolator from 14 dpi to 28 dpi with IBDV (0-14dpi with IAV) to determine IAV shedding. At 17 dpi with IBDV (3 dpi with IAV), six birds from each group were humanely culled to harvest nasal epithelium, trachea, lungs, colon, and kidney in RNAlater (ThermoFisher Scientific) for RNA extraction. The remaining birds were culled at 28 dpi with IBDV (14dpi with IAV) and anti-IAV antibodies in the serum quantified by HI assay (described below).

### Clinical scoring and weights

Birds were weighed daily, and the average daily weight gain calculated. Clinical signs were evaluated as being mild, moderate, or severe based on a clinical scoring system previously used at The Pirbright Institute. If birds exhibiting severe clinical signs, they were promptly and humanely euthanized in adherence to predefined humane endpoints.

### Viral load quantification in tissues

To extract RNA, 750 µl of Trizol LS (Life Technologies) was added to each tissue sample and the mixture was homogenized. Chloroform was added, and the resulting RNA was purified with an RNeasy kit (Qiagen). Reverse transcription quantitative polymerase chain rection (RTqPCR) was performed with a Superscript III Platinum One-Step qRT-PCR Kit (Life Technologies) using primers and a TaqMan probe specific to a conserved region of the influenza A matrix gene, following the method described by Spackman et al. (2002) (49). The thermal cycling conditions were: 50°C for 5 minutes, 95°C for 2 minutes, followed by 40 cycles of 95°C for 3 seconds and 60°C for 30 seconds. Amplification was performed using a 7500 Fast Real-Time PCR System (Applied Biosystems). To create a standard curve, a T7-transcribed RNA standard for the M gene was included in each assay for accurate quantification.

### Hemagglutination-inhibition (HI)

The titer of the anti-IAV antibodies in the serum of the birds was quantified at 14 days post-IAV infection, by HI assay, as previously described (50), using 1% chicken red-blood cells.

### Single cell isolation and flow cytometry

PBMCs were isolated using well-established protocols (33). Briefly, PBMCs were extracted from chicken blood samples by layering onto Histopaque®-1083 Lymphocyte Separation Media (Sigma-Aldrich), and centrifugation at 400 × g for 30 minutes without braking. The purified PBMCs were collected from the interface of the Histopaque and plasma and washed twice with sterile PBS. The staining procedure followed established protocols (33). Briefly, cells were incubated in blocking buffer (4% BSA in PBS) to block Fc receptors, followed by incubation in a cocktail of anti-Bu1-FITC (Cambridge Bioscience) as a marker for B cells, anti-KUL01-PE (Cambridge Bioscience) as a marker for monocytes/macrophages, anti-CD4-PE/Cy7 (Cambridge Bioscience) and anti-CD8α-Pacific Blue (Cambridge Bioscience) as markers for T cells, and Live/Dead™ fixable Near-IR dead cell stain (ThermoFisher). Cells were stained at 4°C for 15 minutes. The cells were then washed twice with fluorescence-activated cell sorter (FACS) buffer (Sigma-Aldrich), fixed with 1% paraformaldehyde at 4°C for 15 minutes, and resuspended in FACS buffer. The cells were then processed using the BD LSRFortessa™ Cell Analyzer (BD Biosciences). Subsequently, data analysis was performed using FlowJo software (v10.8.1) after establishing compensation settings corresponding to monocolour and isotype control stains.

### Titration of infectious virus from swab samples

Sterile polyester-tipped swabs (Fisher Scientific, Loughborough, UK) were used for swabbing of the oropharyngeal and cloacal cavity. Swabs were collected and placed into viral transport media (51), and vortexed. The media was then clarified by centrifugation and stored at −80 °C for subsequent virus detection. The amount of infectious virus in the swab sample liquid was titrated by plaque assay on MDCK cells, following established protocols (48). Briefly, 10-fold serial dilutions of the swab sample liquid was applied to MDCK cells that were subsequently overlaid with 0.6% agarose (Oxoid) in Minimum Essential Medium (MEM), 0.21% bovine serum albumin (BSA), 1 mM l-glutamate, 0.15% sodium bicarbonate, 10 mM HEPES, 1× penicillin/streptomycin (all Gibco), and 0.01% Dextran DEAE (Sigma-Aldrich), along with 2µg/mL TPCK trypsin (Sigma-Aldrich). The cells were then incubated at 37 °C for 72 hours. Plaques were visualized using a crystal violet stain solution containing methanol, and enumerated.

### Sequencing of IAV genomes from the swab samples

The indicated swab samples that were positive for IAV by plaque assay were inoculated into the allantoic cavity of 9-day-old specific pathogen free (SPF) embryonated hen’s eggs (VALO BioMedia GmbH) using standard methods (52). The allantoic fluid was harvested 72 hpi, and viral RNA (vRNA) was extracted using a QIAmp Viral RNA kit (Qiagen), according to the manufacturer’s instructions. The whole IAV genome was amplified from the samples by following previously established protocols (53), with slight modifications. Briefly, cDNA synthesis was initiated by mixing 3 μL of RNA template with Optil-R1 primer (5’-GTT ACG CGC CAG TAG AAA CAA GG-3’) and heating at 65°C for 5 minutes. Subsequently, SuperScript™ IV Reverse Transcriptase (ThermoFisher) was added to achieve a final reaction volume of 20 μl. The reaction proceeded for one cycle at 55°C for 60 minutes, followed by a denaturation step at 80°C for 10 minutes. For PCR amplification of the whole genome, Q5 Hot Start High-Fidelity DNA Polymerase (NEB) was employed, along with 3 μL of cDNA, primers Optil-F1 (5’-GTT ACG CGC CAG CAA AAG CAG G-3’), Optil-F2 (5’-GTT ACG CGC CAGCGA AAG CAG G-3’), and Optil-R1, according to the manufacturer’s recommendations. The PCR cycling conditions were as follows: initial denaturation at 94°C for 2 minutes, followed by 5 cycles of denaturation at 94°C for 30 seconds, annealing at 44°C for 30 seconds, and extension at 68°C for 3.5 minutes. This was succeeded by 26 cycles of denaturation at 94°C for 30 seconds, annealing at 57°C for 30 seconds, and extension at 68°C for 3.5 minutes, with a final extension step at 68°C for 10 minutes. Following PCR amplification, dsDNA samples were diluted to 0.2ng/µL and a total of 1 ng was processed using the Nextera XT DNA library preparation kit as per the manufacturers protocols. This was automated using a Hamilton NGStar (Hamilton Robotics). Libraries were bead normalized according to manufacturers protocols and pooled for sequencing. Sequencing pools were spiked with 1% PhiX and run on an Illumina MiSeq with a 2x250 cycle v2 reagent kit. The sequencing data was aligned to a reference genome of all 8 segments of the A/Mallard/Alberta/156/01 (H3N8) strain (Accession numbers CY004702.1- CY004709.1) (23) with BWA-mem version 0.7.17-r1188. The variants were called from the resulting alignment using freebayes version v0.9.21. These variants were annotated with VEP version 107.0 using a custom database to show amino acid changes and their consequences. To calculate the pi-diversity, the pileup of the alignments was run through bcftools (version 1.10.2) multi-allele caller, and the result was piped to vcftools (version 0.1.16) with the –site-pi option.

The stock of H3N8 that we used to inoculate the birds (the inoculum) was also sequenced in the same way and compared to the reference genome.

### Structural modeling

The amino acid sequences of the HA proteins from the inoculum and from the swab samples with the specified mutations were used to model the structure of the HA homotrimers, and the amino acid sequences of the PB2, PB1, and PA from the inoculum and from the swab samples with the specified mutations were used to model the structure of the heterohexamer polymerase complexes. The structures were modelled using AlphaFold3 through the AlphaFold Server platform by modeling three instances (HA) or two instances (each of PB2, PB1, and PA) of a protein entity bearing the specified mutations. Selection of a single PB2/PB1/PA trimer and rendering of all models was performed with UCSF Chimera X.

### Statistical analyses

Average daily weight gains, immune cell populations quantified by flow cytometry, and viral replication quantified by RTqPCR and plaque assay were compared by a one-way analysis of variance (ANOVA) with Tukey *post hoc* comparisons using GraphPad Prism version 10.0 (GraphPad Software, Inc., San Diego, CA). The amino acid substitutions per sample were compared between Mock/IAV and IBDV/IAV groups by an unpaired Students t-test. Results were considered significantly different when *p* was <0.05.

## Acknowledgements

This work was supported by grants BB/T008806/1, BBS/E/I/00007030, BBS/E/I/00007035, BBS/E/I/00007039, and BBS/E/PI/230002C, funded by Biotechnology and Biological Sciences Research Council, U.K. For the purpose of Open Access, the author has applied a CC BY public copyright license to any Author Accepted Manuscript version arising from this submission. The funders had no role in study design, data collection and interpretation, or the decision to submit the work for publication.

## Supporting Information

**S1 Table. H3N8 IAV nucleotide variants and amino acid substitutions detected in individual swab samples.** The nucleotide variants that were detected in the H3N8 IAV oropharyngeal swab samples from Mock/IAV and IBDV/IAV groups are listed, together with their location in the viral gene segment, the frequency of the reads in which the variant was detected, and the read depth. Where the nucleotide variant led to an amino acid substitution, the location of the substitution in the gene segment and the lettered amino acid substitution is listed. Entries highlighted in red had a read depth of 1,000 or over.

## References

1. Wille M, Latorre-Margalef N, Tolf C, Halpin R, Wentworth D, Fouchier RAM, et al. Where do all the subtypes go? Temporal dynamics of H8-H12 influenza A viruses in waterfowl. Virus Evol. 2018;4(2):vey025.

2. Carnaccini S, Perez DR. H9 Influenza Viruses: An Emerging Challenge. Cold Spring Harb Perspect Med. 2020;10(6).

3. Everest H, Billington E, Daines R, Burman A, Iqbal M. The Emergence and Zoonotic Transmission of H10Nx Avian Influenza Virus Infections. mBio. 2021;12(5):e0178521.

4. Everest H, Hill SC, Daines R, Sealy JE, James J, Hansen R, et al. The Evolution, Spread and Global Threat of H6Nx Avian Influenza Viruses. Viruses. 2020;12(6).

5. Jimenez-Bluhm P, Karlsson EA, Ciuoderis KA, Cortez V, Marvin SA, Hamilton-West C, et al. Avian H11 influenza virus isolated from domestic poultry in a Colombian live animal market. Emerg Microbes Infect. 2016;5(12):e121.

6. Krauss S, Stucker KM, Schobel SA, Danner A, Friedman K, Knowles JP, et al. Long-term surveillance of H7 influenza viruses in American wild aquatic birds: are the H7N3 influenza viruses in wild birds the precursors of highly pathogenic strains in domestic poultry? Emerg Microbes Infect. 2015;4(6):e35.

7. Lee DH, Torchetti MK, Hicks J, Killian ML, Bahl J, Pantin-Jackwood M, et al. Transmission Dynamics of Highly Pathogenic Avian Influenza Virus A(H5Nx) Clade 2.3.4.4, North America, 2014-2015. Emerg Infect Dis. 2018;24(10):1840-8.

8. Lin S, Zhang Y, Yang J, Yang L, Li X, Bo H, et al. Cross-Species Transmission Potential of H4 Avian Influenza Viruses in China: Epidemiological and Evolutionary Study. Viruses. 2024;16(3).

9. Mo J, Youk S, Pantin-Jackwood MJ, Suarez DL, Lee DH, Killian ML, et al. The pathogenicity and transmission of live bird market H2N2 avian influenza viruses in chickens, Pekin ducks, and guinea fowl. Vet Microbiol. 2021;260:109180.

10. Yang J, Zhang Y, Yang L, Li X, Bo H, Liu J, et al. Evolution of Avian Influenza Virus (H3) with Spillover into Humans, China. Emerg Infect Dis. 2023;29(6):1191–201.

11. Pepin KM, Spackman E, Brown JD, Pabilonia KL, Garber LP, Weaver JT, et al. Using quantitative disease dynamics as a tool for guiding response to avian influenza in poultry in the United States of America. Prev Vet Med. 2014;113(4):376–97.

12. Peacock T, Moncla L, Dudas G, VanInsberghe D, Sukhova K, Lloyd-Smith JO, et al. The global H5N1 influenza panzootic in mammals. Nature. 2024.

13. Hoerr FJ. Clinical aspects of immunosuppression in poultry. Avian Dis. 2010;54(1):2–15.

14. Ingrao F, Rauw F, Lambrecht B, van den Berg T. Infectious Bursal Disease: a complex host-pathogen interaction. Dev Comp Immunol. 2013;41(3):429–38.

15. van den Berg TP, Eterradossi N, Toquin D, Meulemans G. Infectious bursal disease (Gumboro disease). Rev Sci Tech. 2000;19(2):509–43.

16. Chaudhry M, Rashid HB, Thrusfield M, Welburn S, Bronsvoort BM. A case-control study to identify risk factors associated with avian influenza subtype H9N2 on commercial poultry farms in Pakistan. PLoS One. 2015;10(3):e0119019.

17. Spackman E, Stephens CB, Pantin-Jackwood MJ. The Effect of Infectious Bursal Disease Virus-Induced Immunosuppression on Vaccination Against Highly Pathogenic Avian Influenza Virus. Avian Dis. 2018;62(1):36–44.

18. Khalil NW, Elshorbagy MA, Elboraay EM, Helal AM. Live IBD vaccine exacerbates disease and pathological effects of Asian lineage H9N2 LPAIV in chickens. Avian Pathol. 2023;52(5):351–61.

19. Ranjbar VR, Mohammadi A, Dadras H. Infectious bursal disease virus suppresses H9N2 avian influenza viral shedding in broiler chickens. Br Poult Sci. 2019;60(5):493–8.

20. Hashemzade F, Mayahi M, Shoshtary A, Reza Seyfi Abad Shapouri M, Ghorbanpoor M. Effect of experimental infectious bursal disease virus on clinical signs and pathogenesis of avian influenza virus H(9)N(2) in turkey by real time PCR. Vet Res Forum. 2019;10(4):293-7.

21. Ramirez-Nieto G, Shivaprasad HL, Kim CH, Lillehoj HS, Song H, Osorio IG, et al. Adaptation of a mallard H5N2 low pathogenicity influenza virus in chickens with prior history of infection with infectious bursal disease virus. Avian Dis. 2010;54(1 Suppl):513-21.

22. Nazki S, Reddy V, Kamble N, Sadeyen JR, Iqbal M, Behboudi S, et al. CD4(+)TGFbeta(+) cells infiltrated the bursa of Fabricius following IBDV infection, and correlated with a delayed viral clearance, but did not correlate with disease severity, or immunosuppression. Front Immunol. 2023;14:1197746.

23. Obenauer JC, Denson J, Mehta PK, Su X, Mukatira S, Finkelstein DB, et al. Large-scale sequence analysis of avian influenza isolates. Science. 2006;311(5767):1576-80.

24. Bahl J, Krauss S, Kuhnert D, Fourment M, Raven G, Pryor SP, et al. Influenza a virus migration and persistence in North American wild birds. PLoS Pathog. 2013;9(8):e1003570.

25. Courtillon C, Allee C, Amelot M, Keita A, Bougeard S, Hartle S, et al. Blood B Cell Depletion Reflects Immunosuppression Induced by Live-Attenuated Infectious Bursal Disease Vaccines. Front Vet Sci. 2022;9:871549.

26. Bertran K, Lee DH, Criado MF, Smith D, Swayne DE, Pantin-Jackwood MJ. Pathobiology of Tennessee 2017 H7N9 low and high pathogenicity avian influenza viruses in commercial broiler breeders and specific pathogen free layer chickens. Vet Res. 2018;49(1):82.

27. James J, Billington E, Warren CJ, De Sliva D, Di Genova C, Airey M, et al. Clade 2.3.4.4b H5N1 high pathogenicity avian influenza virus (HPAIV) from the 2021/22 epizootic is highly duck adapted and poorly adapted to chickens. J Gen Virol. 2023;104(5).

28. Kwon JH, Bertran K, Lee DH, Criado MF, Killmaster L, Pantin-Jackwood MJ, et al. Diverse infectivity, transmissibility, and pathobiology of clade 2.3.4.4 H5Nx highly pathogenic avian influenza viruses in chickens. Emerg Microbes Infect. 2023;12(1):2218945.

29. Leyson C, Youk SS, Smith D, Dimitrov K, Lee DH, Larsen LE, et al. Pathogenicity and genomic changes of a 2016 European H5N8 highly pathogenic avian influenza virus (clade 2.3.4.4) in experimentally infected mallards and chickens. Virology. 2019;537:172-85.

30. Pantin-Jackwood MJ, Stephens CB, Bertran K, Swayne DE, Spackman E. The pathogenesis of H7N8 low and highly pathogenic avian influenza viruses from the United States 2016 outbreak in chickens, turkeys and mallards. PLoS One. 2017;12(5):e0177265.

31. Chrzastek K, Segovia K, Torchetti M, Killian ML, Pantin-Jackwood M, Kapczynski DR. Virus Adaptation Following Experimental Infection of Chickens with a Domestic Duck Low Pathogenic Avian Influenza Isolate from the 2017 USA H7N9 Outbreak Identifies Polymorphic Mutations in Multiple Gene Segments. Viruses. 2021;13(6).

32. Asfor AS, Nazki S, Reddy V, Campbell E, Dulwich KL, Giotis ES, et al. Transcriptomic Analysis of Inbred Chicken Lines Reveals Infectious Bursal Disease Severity Is Associated with Greater Bursal Inflammation In Vivo and More Rapid Induction of Pro-Inflammatory Responses in Primary Bursal Cells Stimulated Ex Vivo. Viruses. 2021;13(5).

33. Nazki S, Reddy VRAP, Kamble N, Sadeyen J-R, Iqbal M, Behboudi S, et al. CD4+TGFβ+ cells infiltrated the bursa of Fabricius following IBDV infection, and correlated with a delayed viral clearance, but did not correlate with disease severity, or immunosuppression. Frontiers in Immunology. 2023;14.

34. Ferreri LM, Geiger G, Seibert B, Obadan A, Rajao D, Lowen AC, et al. Intra- and inter-host evolution of H9N2 influenza A virus in Japanese quail. Virus Evol. 2022;8(1):veac001.

35. Youk SS, Leyson CM, Seibert BA, Jadhao S, Perez DR, Suarez DL, et al. Mutations in PB1, NP, HA, and NA Contribute to Increased Virus Fitness of H5N2 Highly Pathogenic Avian Influenza Virus Clade 2.3.4.4 in Chickens. J Virol. 2021;95(5).

36. Hiromoto Y, Saito T, Lindstrom S, Nerome K. Characterization of low virulent strains of highly pathogenic A/Hong Kong/156/97 (H5N1) virus in mice after passage in embryonated hens’ eggs. Virology. 2000;272(2):429–37.

37. Fu Y, Wedde M, Smola S, Oh DY, Pfuhl T, Rissland J, et al. Different populations of A(H1N1)pdm09 viruses in a patient with hemolytic-uremic syndrome. Int J Med Microbiol. 2024;314:151598.

38. Burke DF, Smith DJ. A recommended numbering scheme for influenza A HA subtypes. PLoS One. 2014;9(11):e112302.

39. Legnardi M, Franzo G, Tucciarone CM, Koutoulis K, Cecchinato M. Infectious bursal disease virus in Western Europe: the rise of reassortant strains as the dominant field threat. Avian Pathol. 2023;52(1):25–35.

40. Reddy V, Bianco C, Poulos C, Egana-Labrin SC, Brodrick AJ, Nazki S, et al. Molecular characterization of reassortant infectious bursal disease virus (IBDV) strains of genogroup A3B1 detected in some areas of Britain between 2020 and 2021. Virology. 2024;600:110269.

41. Michel LO, Jackwood DJ. Classification of infectious bursal disease virus into genogroups. Arch Virol. 2017;162(12):3661–70.

42. Reddy V, Nazki S, Brodrick AJ, Asfor A, Urbaniec J, Morris Y, et al. Evaluating the Breadth of Neutralizing Antibody Responses Elicited by Infectious Bursal Disease Virus Genogroup A1 Strains Using a Novel Chicken B-Cell Rescue System and Neutralization Assay. J Virol. 2022;96(18):e0125522.

43. Zhang W, Wang X, Gao Y, Qi X. The Over-40-Years-Epidemic of Infectious Bursal Disease Virus in China. Viruses. 2022;14(10).

44. Baba TW, Giroir BP, Humphries EH. Cell lines derived from avian lymphomas exhibit two distinct phenotypes. Virology. 1985;144(1):139–51.

45. Bayliss CD, Spies U, Shaw K, Peters RW, Papageorgiou A, Muller H, et al. A comparison of the sequences of segment A of four infectious bursal disease virus strains and identification of a variable region in VP2. J Gen Virol. 1990;71 ( Pt 6):1303–12.

46. Dulwich KL, Asfor A, Gray A, Giotis ES, Skinner MA, Broadbent AJ. The Stronger Downregulation of in vitro and in vivo Innate Antiviral Responses by a Very Virulent Strain of Infectious Bursal Disease Virus (IBDV), Compared to a Classical Strain, Is Mediated, in Part, by the VP4 Protein. Front Cell Infect Microbiol. 2020;10:315.

47. Reed LJ, Muench H. A simple method of estimating fifty percent end points. Am J Hyg. 1938;27:493–7.

48. James J, Howard W, Iqbal M, Nair VK, Barclay WS, Shelton H. Influenza A virus PB1-F2 protein prolongs viral shedding in chickens lengthening the transmission window. J Gen Virol. 2016;97(10):2516–27.

49. Spackman E, Senne DA, Myers TJ, Bulaga LL, Garber LP, Perdue ML, et al. Development of a real-time reverse transcriptase PCR assay for type A influenza virus and the avian H5 and H7 hemagglutinin subtypes. J Clin Microbiol. 2002;40(9):3256–60.

50. Broadbent AJ, Santos CP, Anafu A, Wimmer E, Mueller S, Subbarao K. Evaluation of the attenuation, immunogenicity, and efficacy of a live virus vaccine generated by codon-pair bias de-optimization of the 2009 pandemic H1N1 influenza virus, in ferrets. Vaccine. 2016;34(4):563–70.

51. Richard M, van den Brand JMA, Bestebroer TM, Lexmond P, de Meulder D, Fouchier RAM, et al. Influenza A viruses are transmitted via the air from the nasal respiratory epithelium of ferrets. Nat Commun. 2020;11(1):766.

52. Senne D. Virus propagation in embryonating eggs. A laboratory manual for the isolation and identification of avian pathogens. 1998:235–40.

53. Chrzastek K, Lee DH, Gharaibeh S, Zsak A, Kapczynski DR. Characterization of H9N2 avian influenza viruses from the Middle East demonstrates heterogeneity at amino acid position 226 in the hemagglutinin and potential for transmission to mammals. Virology. 2018;518:195–201.

